# A High-Throughput, Flow Cytometry Approach to Measure Phase Behavior and Exchange in Biomolecular Condensates

**DOI:** 10.1101/2025.06.02.657082

**Authors:** Yuchen He, George M. Ongwae, Anupam Mondal, Jeetain Mittal, Marcos M. Pires

## Abstract

Biomolecular condensates are essential for cellular organization, yet their formation dynamics and molecular content exchange properties remain poorly understood. In this study, we developed a high-throughput flow cytometry approach to quantify condensate formation, molecular colocalization, and dynamic exchange. Using self-interacting NPM1 condensates as a model system, we benchmarked the use of flow cytometry to broadly characterize their phase behavior across various protein concentrations and salt conditions. We further demonstrated that flow cytometry can assess the colocalization of macromolecules - including antibodies, lipids, small-molecule drugs, and RNAs, within NPM1 condensates. Importantly, we established the first assay to track real-time molecular exchange between preformed condensates and newly added, orthogonally tagged protein. This revealed that condensate aging significantly reduces molecular dynamisms, likely due to altered biophysical properties with time. Compared to conventional imaging techniques, which often require immobilized samples and complex experimental setups, our solution-based approach enables rapid, quantitative analysis of condensate behavior at the single-droplet level. This work provides a powerful framework for studying biomolecular condensates with enhanced precision and scalability, offering new insights into their dynamic properties and molecular interactions.

## INTRODUCTION

Biomolecular condensates represent a fundamental organizing principle in cell biology, enabling the compartmentalization and regulation of biochemical reactions without the need for membrane-bound organelles.^1, 2^ These dynamic structures form via liquid-liquid phase separation (LLPS), a physicochemical process in which biomolecules demix from the surrounding milieu to generate dense, droplet-like assemblies.^3, 4^ LLPS is driven by multivalent interactions among proteins and nucleic acids, often facilitated by intrinsically disordered regions (IDRs), low-complexity domains, and specific modular interaction motifs.^5–8^ By concentrating proteins, nucleic acids, and other macromolecules, condensates orchestrate important biochemical and biophysical processes such as gene regulation, signal transduction, RNA metabolism, and stress response (**Fig. 1a**).^9–11^ Their ability to assemble and disassemble in response to environmental cues is essential for cellular homeostasis and maintenance. Disruptions in condensate formation or changes in their material properties have been implicated in various pathologies including amyotrophic lateral sclerosis (ALS), frontotemporal dementia, and several cancers.^12–16^ Understanding the molecular principles governing condensate assembly, composition, and dynamics is therefore critical for elucidating cellular functional organization and developing targeted therapies.

**Fig. 1.**
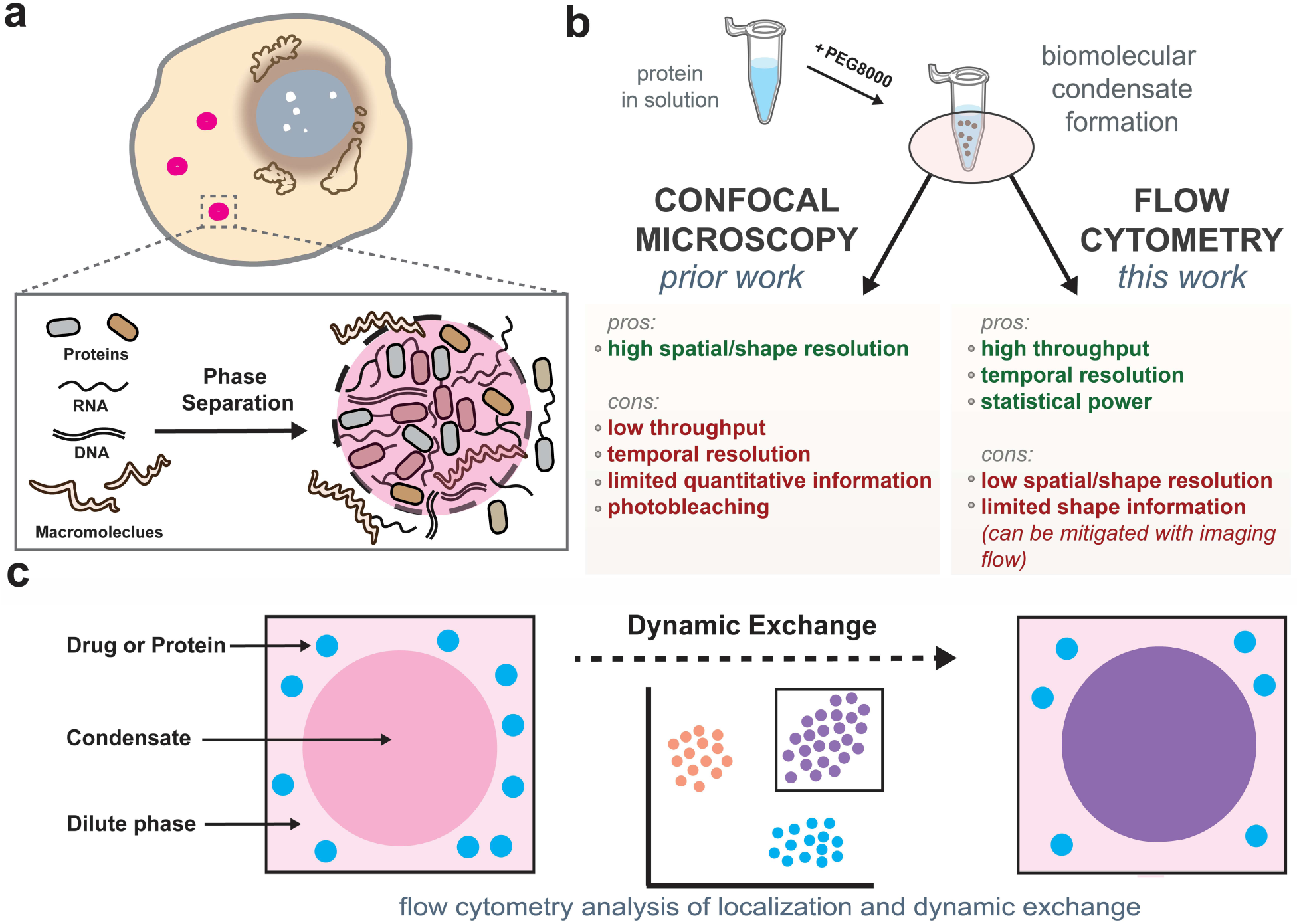
Developing flow cytometry-based strategies to study biomolecular condensates. **a)** Schematic illustration of biomolecular condensate formation via liquid– liquid phase separation (LLPS) from proteins, RNAs, and DNAs in cells.z **b)** Comparison of confocal microscopy and flow cytometry for condensate analysis. **c)** Conceptual overview showing dynamic exchange between condensates and surrounding molecules. Flow cytometry enables continuous tracking of such exchange events over time, providing a powerful approach to assess condensate dynamics.

As our understanding of condensate biology expands, increasing attention has turned to how specific partner molecules interact with or partition into these structures.^17–19^ While initial studies largely examined the roles of proteins and nucleic acids in driving or enriching within condensates, recent work has broadened this view to include small molecules. For example, one early discovery revealed that cyclic GMP-AMP synthase (cGAS) can undergo phase separation upon binding to oligonucleotides, with these nucleic acids enriching within the resulting condensates.^20^ Building on this, researchers have begun to systematically investigate which classes of molecules can diffuse into, concentrate within, or be excluded from existing condensates. These efforts have primarily focused on small molecules, particularly in the context of how condensates might influence drug distribution within cells. Notably, the Young lab demonstrated that fluorophore-tagged FDA-approved therapeutics showed differential localization across diverse protein condensates.^21^ This work has since expanded to BODIPY-based libraries to further explore physicochemical determinants of condensate partitioning.^22^ More recently, a wide scan of over 1000 small-molecule drugs were evaluated for their colocalization with condensates.^23^

Despite growing recognition of their significance, *in vitro* analyses of biomolecular condensates remain challenging tasks, largely constrained by methodological limitations. Confocal microscopy has long been the primary method used to analyze condensate formation, dynamics, and composition, offering high spatial resolution.^20–22^ However, its low throughput limits experimental scalability, and the temporal resolution is often insufficient to capture rapid dynamics. Manual image processing can also introduce user bias, reducing statistical confidence. Other commonly used methods, such as fluorescence recovery after photobleaching (FRAP), turbidity measurement, and sedimentation assays,^24–26^ provide insights into molecular mobility or condensate composition but are either low throughput or lack spatial and temporal resolution. These techniques often fail to capture condensate heterogeneity or rapid dynamic changes in a quantitative manner. While alternative tools such as microfluidic platforms are being explored,^27^ there remains a need for robust, quantitative, and scalable methods to study condensate behavior.

To address these limitations, we provide direct evidence that flow cytometry can be a superior method for mimicking freedom of condensate in a medium towards high-throughput and quantitative analysis of biomolecular condensates (**Fig. 1b**). Flow cytometry offers superior quantitative analysis of protein condensates compared to confocal microscopy through its ability to measure thousands to millions of individual droplets within minutes, providing robust statistical power to detect condensate heterogeneity across droplet pools. Unlike confocal microscopy, which can be limited by photobleaching, variable illumination, and smaller sample sizes, flow cytometry delivers highly reproducible fluorescence measurements with superior signal-to-noise ratios under consistent excitation conditions. This method can propel the field forward in terms of real-time kinetic analysis of condensate formation and dissolution, capturing rapid dynamic processes with high temporal resolution while minimizing artifactual effects. This high-throughput capability makes flow cytometry particularly valuable for screening applications, enabling systematic evaluation of condensate modulation across multiple experimental conditions simultaneously, ultimately providing more statistically robust and physiologically relevant measurements of condensate properties than traditional confocal approaches.

Our data shows that flow cytometry reliably captures condensate formation, composition, and dissolution, closely aligning with imaging-based observations. Importantly, this technique also enables real-time monitoring of protein exchange between condensates and their surroundings (**Fig. 1c**), offering new kinetic insights into phase-separated systems. To further dissect the molecular basis of condensate formation, we complemented our experimental work with coarse-grained molecular dynamics simulations. In parallel, to mechanistically understand the exchange dynamics during aging, we developed a discrete-state stochastic model that quantitatively supports our experimental findings using first-passage calculations. By expanding the analytical toolkit for condensate research, this study establishes flow cytometry as a robust and scalable method for probing phase-separated systems. Its ability to resolve dynamic behaviors with high temporal and statistical precision offers new opportunities for mechanistic discovery, with potential implications for therapeutic targeting of condensate-associated pathologies.

## RESULTS AND DISCUSSIONS

### Characterization of condensate formation and dissolution

Unlike cellular condensates that assemble under physiological conditions, *in vitro* condensate formation typically requires crowding agents to drive phase separation. Reagents such as polyethylene glycol (PEG) and Ficoll mimic the intracellular macromolecular environment by increasing effective protein concentration and promoting demixing through excluded volume effects.^28, 29^ As an initial test, we selected four well-characterized scaffolding proteins — NPM1, HP1α, DDX4, and HMGB1 — each previously reported to undergo phase separation and function in chromatin organization, RNA metabolism, and nucleolar assembly.^30–34^ Each protein was fused to a fluorescent tag to facilitate visualization of condensate behavior (**Fig. 2a**). The recombinant proteins were purified and subjected to *in vitro* droplet assays to assess condensate formation. Specifically, each protein was incubated with 10% (w/v) PEG8000 for 30 minutes and initially imaged using confocal microscopy as previously described (**Fig. 2a**). To assess the sensitivity of these condensates to disruption, we treated the assembled droplets with 1,6-hexanediol, a known disruptor of weak hydrophobic and hydrogen-bonding interactions in biomolecular condensates.^35, 36^ As expected, each protein formed spherical condensates in the presence of PEG8000, confirming their phase separation propensity (**Fig. 2b, e-g**). Addition of 1,6-hexanediol led to the dissolution of these structures, demonstrating the reversibility of condensate formation.

**Figure 2.**
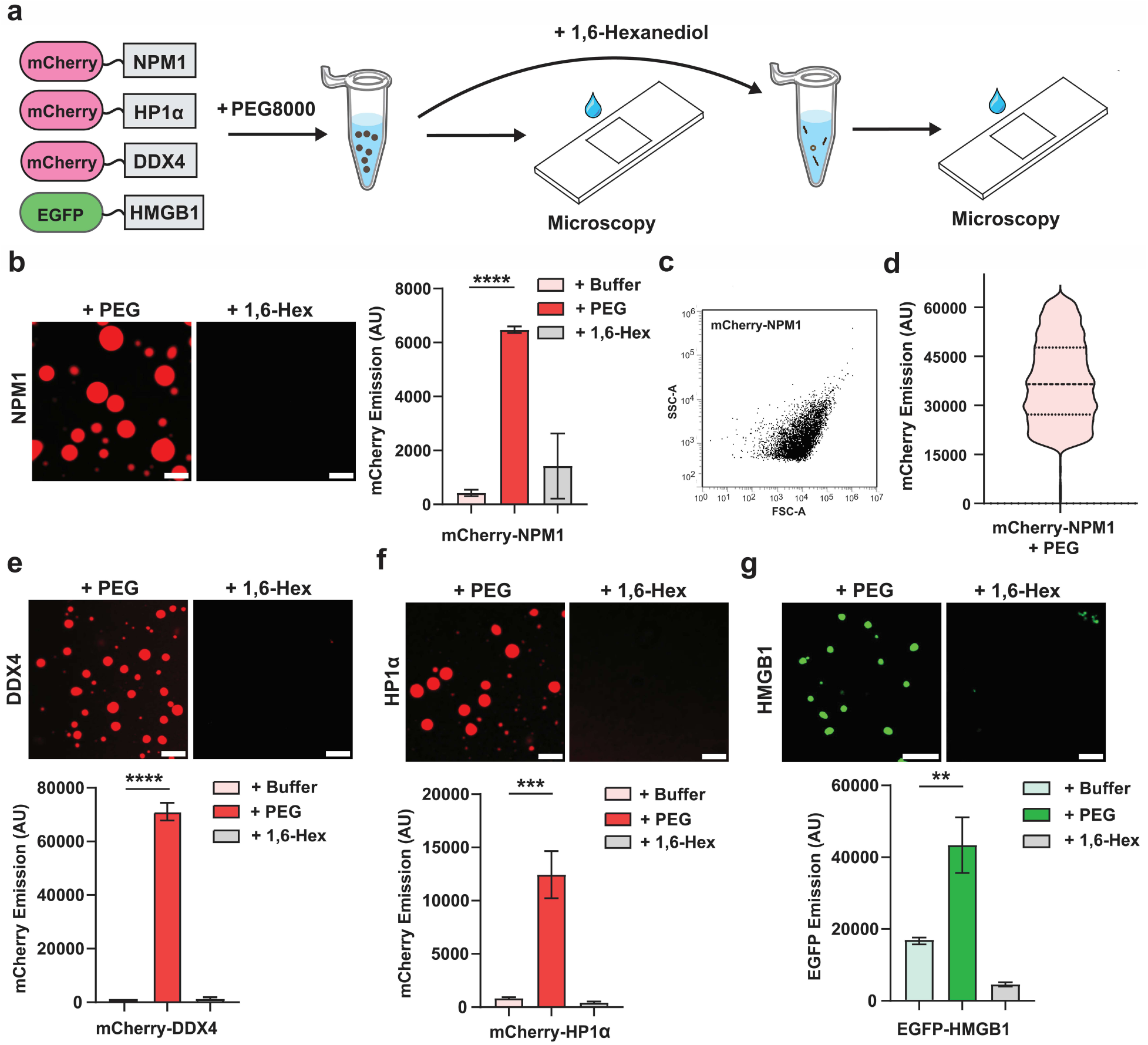
Characterization of condensate formation and dissolution using confocal microscopy and flow cytometry. **a)** Schematic overview of the *in vitro* droplet assay workflow. Purified fluorescently tagged proteins (mCherry-NPM1, mCherry-DDX4, mCherry-HP1α, and EGFP-HMGB1) were incubated with 10% PEG8000 to induce condensate formation, followed by treatment with 10% 1,6-hexanediol to assess reversibility. **b)** Confocal images and flow cytometry analysis of 20 µM mCherry-NPM1 under control, PEG, and PEG + hexanediol conditions. PEG treatment induces bright, spherical condensates, which are disrupted by 1,6-hexanediol. Flow cytometry confirms increased fluorescence intensity with PEG and reduction upon hexanediol treatment, indicating reversible condensate formation. **c)** Representative side scatter (SSC-A) vs. forward scatter (FSC-A) plot of 20 µM mCherry-NPM1 in the presence of PEG8000, highlighting PEG-induced shifts in condensate size and granularity. **d)** Violin plot showing the distribution of fluorescence intensities for 5,000 gated mCherry-NPM1 condensates. The central dashed line represents the median, and the outer dashed lines indicate the first and third quartiles. **e-g)** Confocal microscopy and flow cytometry analyses of mCherry-DDX4 (**e**), mCherry-HP1α (**f**), and EGFP-HMGB1 (**g**) under control, PEG, and PEG + hexanediol conditions. Flow cytometry data represents gated populations of fluorescent particles. All datasets are representative of at least three independent replicates. Statistical analysis was performed using unpaired, two-tailed student’s *t*-tests. Error bars indicate standard deviation (SD). Significance is denoted as follows: p < 0.05 (*), p < 0.01 (**), p < 0.001 (***), and p < 0.0001 (****). Scale bars = 5 µm.

To validate these observations using our flow cytometry approach, we analyzed the same samples using flow cytometry. In the presence of PEG8000, a significant increase in mean fluorescence intensity was observed, indicating robust condensate formation and this is expected to have a higher density of fluorescent proteins per event analyzed. Treatment with 1,6-hexanediol led to a marked decrease in fluorescence signal, reflecting condensate dissolution (**Fig. 2b, e-g**). Each condition was analyzed across 10,000 individual events (putative condensates), and all measurements were performed in triplicate to ensure reproducibility. Importantly, flow cytometry also offers biophysical insights beyond fluorescence intensity. For example, side scatter (SSC) and forward scatter (FSC) plots provide estimates of granularity and size distribution. We show this using mCherry-NPM1 as a representative case: PEG treatment causes a clear shift in SSC-A versus FSC-A profiles, consistent with increased size and complexity (**Fig. 2c**). Furthermore, a violin plot of mCherry-NPM1 particle intensities (**Fig. 2d**) illustrates how flow cytometry captures population-level heterogeneity across tens of thousands of events — a resolution that is difficult to achieve by microscopy alone. Scatter and fluorescence distribution plots for the other proteins are shown in **Fig. S1**. These data establish that flow cytometry alone reliably captures condensate formation and dissolution in a high-throughput, statistically robust manner, consistently mirroring the results obtained by conventional imaging. We envision that this modality of analysis could be paired with high volume small molecule libraries to discover novel drug-like agents that can block formation or disrupt droplets.

### Characterization of NPM1 condensates using experimental and computational approaches

To gain deeper mechanistic insights into condensate behavior, we selected one representative scaffold protein, NPM1 (mCherry-NPM1), for detailed characterization. Turbidity measurements of mCherry-NPM1 in the presence of PEG8000 revealed a concentration-dependent increase in light scattering, consistent with enhanced condensate formation (**Fig. 3a**). Real-time monitoring NPM1 further showed rapid increase in turbidity upon PEG addition, indicating fast condensate assembly (**Fig. 3b**). Confocal microscopy revealed that increasing protein concentrations led to the formation of a higher density and visibly larger droplets (**Fig. 3c**), reflecting enhanced phase separation under higher protein concentrations. Parallel flow cytometry analysis showed an analogous increase in mCherry fluorescence intensity with increases in protein concentration (**Fig. 3d**), thus mirroring what was observed using confocal microscopy and consistent with a larger association of fluorescently tagged proteins per event. In addition, forward and side scatter plots (FSC vs. SSC) indicated a size-dependent shift in condensate populations, supporting the expected increase in the size of the formed droplets at higher protein concentrations (**Fig. 3e**).

**Fig. 3.**
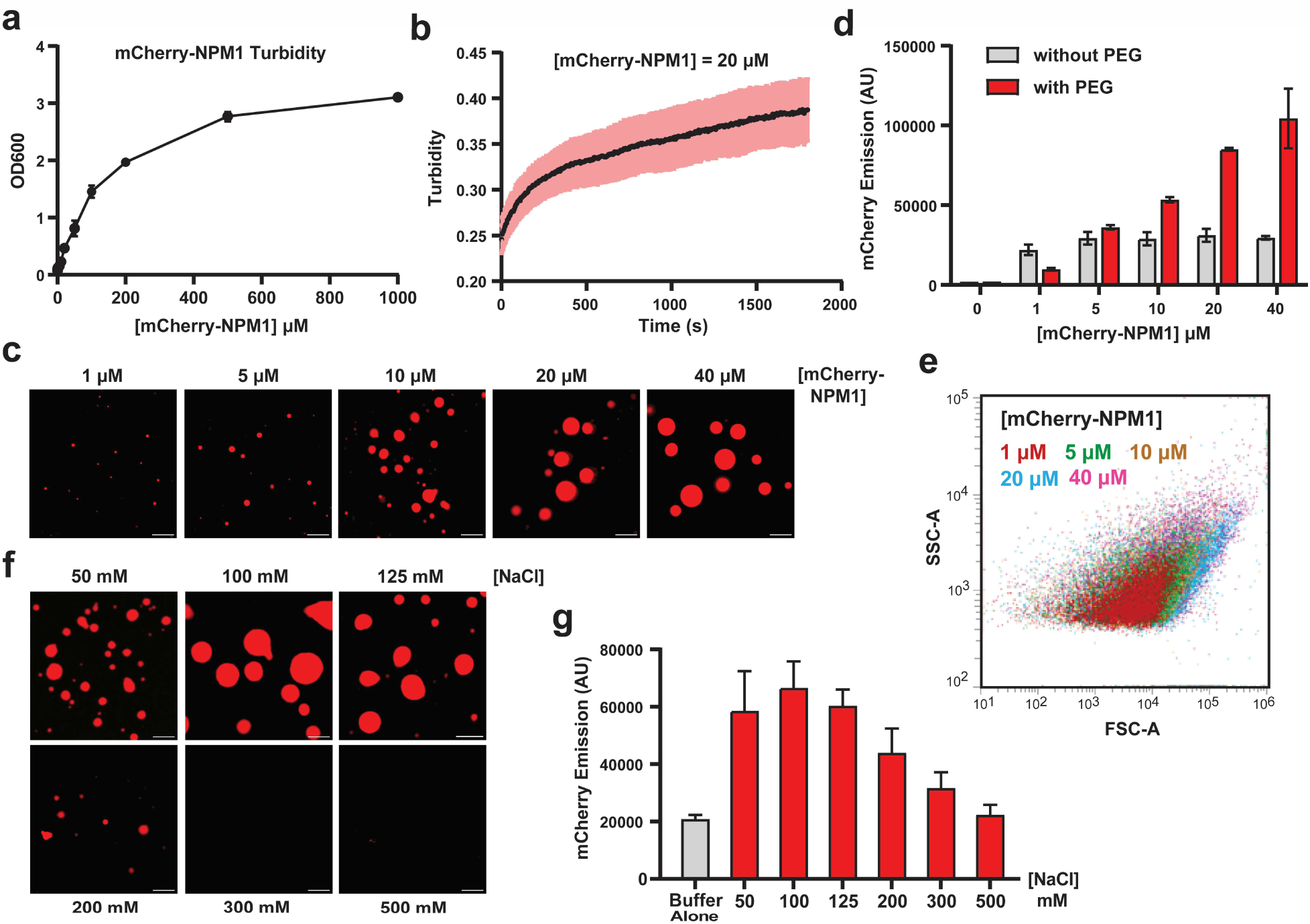
Characterization of NPM1 condensates using a combination of diverse approaches. **a)** Turbidity measurements at OD_600_ show a concentration-dependent increase, indicating higher condensate formation at increasing NPM1 concentrations. **b)** Time-course turbidity measurements of 20 µM NPM1 reveal rapid condensate formation. **b)** Confocal images of mCherry-NPM1 droplets at different protein concentrations show larger droplet sizes with increasing concentration. **d)** Flow cytometry analysis is consistent with **c)**, showing that higher mCherry-NPM1 concentrations result in increased fluorescence intensity, reflecting more condensate formation. **e)** Forward scatter (FSC-A) and side scatter (SSC-A) analysis of mCherry-NPM1 condensates indicate that larger condensates exhibit higher light scattering, consistent with increased droplet size. **f)** Confocal images of 20 µM mCherry-NPM1 droplets in buffers with varying NaCl concentrations show that condensates remain stable below 125 mM NaCl but dissolve at higher salt concentrations (>300 mM). **g)** Flow cytometry analysis aligns with the salt titration data in **f)**, confirming the salt-dependent dissolution of NPM1 condensates. Scale bars = 5 µm.

Given the electrostatic nature of many condensate-driving interactions,^37, 38^ we next benchmarked whether flow cytometry could report on salt sensitivity as had been previously described.^39^ NaCl titration from 50 mM to 500 mM revealed a salt-dependent modulation of condensate stability: condensates remained visible between 50 to 125 mM NaCl but were completely disrupted at 300-500 mM NaCl (**Fig. 3f**). This observation is consistent with electrostatic screening effects, where increased ionic strength weakens multivalent protein-protein and protein-RNA interactions essential for condensate stability.^40^ Flow cytometry results mirrored this trend with corresponding decreases in fluorescence intensity (**Fig. 3g**).

One potential concern with flow cytometry is whether shear stress or buffer conditions may compromise condensate integrity. To address this, we analyzed mCherry-NPM1 condensates using imaging flow cytometry (IFC), which combines the high-throughput, quantitative capabilities of flow cytometry with high-resolution imaging across multiple channels.^41, 42^ While typically used for cellular analysis, IFC has been applied to study protein aggregates and extracellular vesicles,^43, 44^ demonstrating its utility for analyzing smaller particles. IFC confirmed that increasing protein concentrations resulted in larger, brighter condensates visible in both bright field and fluorescence channels (**Fig. 4a**). Plotting surface area (bright field) against fluorescence intensity (mCherry) demonstrated a concentration-dependent shift toward larger, brighter events (**Fig. 4b**). Quantification revealed a concentration-dependent increase in condensate size and fluorescence intensity (**Fig. 4b-c**), consistent with results from confocal microscopy (**Fig. 3c**). The mean fluorescence intensity measured by IFC also closely matched the data obtained from standard flow cytometry (**Fig. 3d**, additional representative images are shown in **Fig. S2**). Critically, these results provided key evidence that the events registered in flow mirror the general dimensional features of the condensates observed using standard confocal microscopy.

**Fig. 4.**
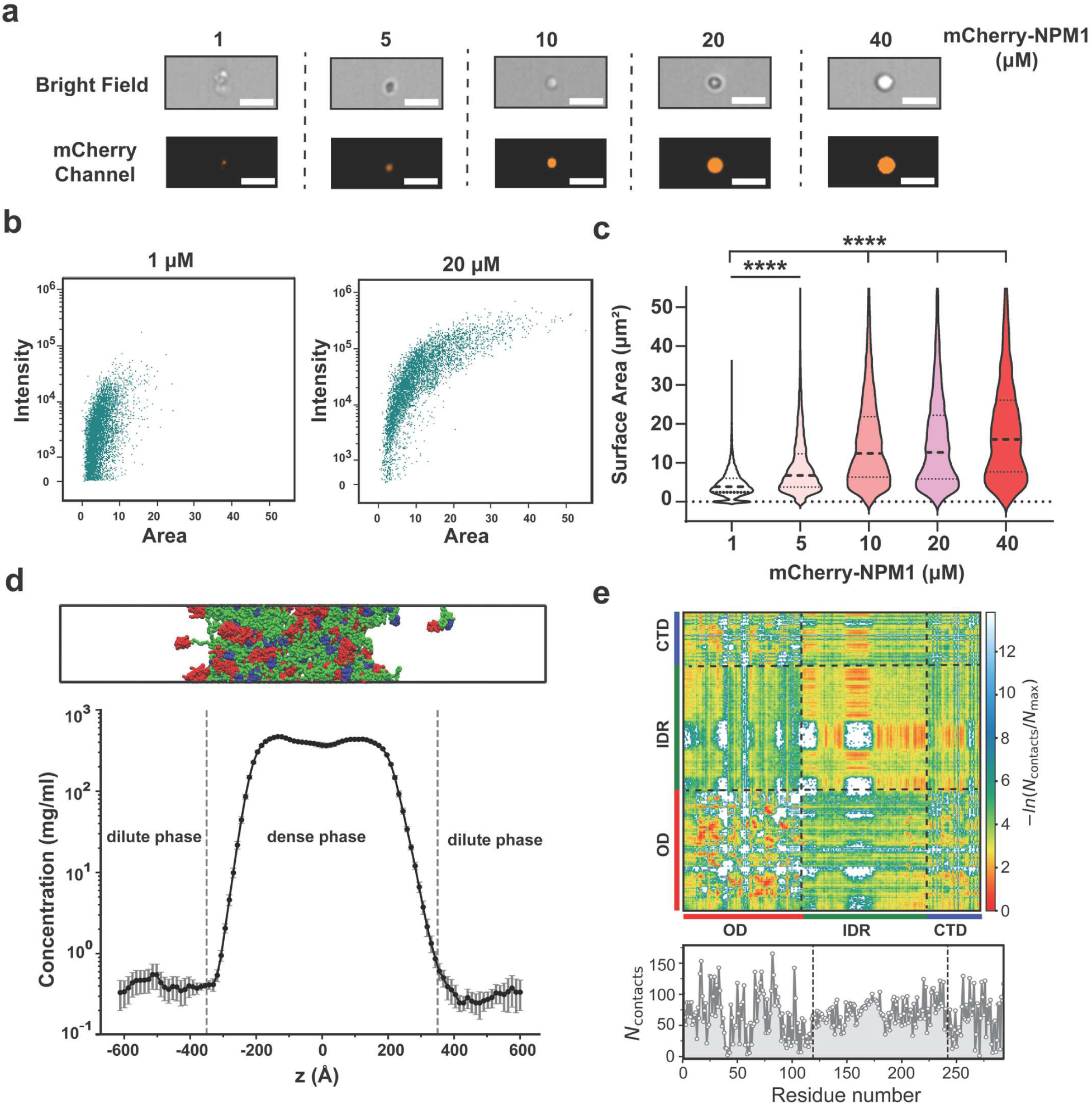
Imaging flow cytometry (IFC) analysis of mCherry-NPM1 condensates and characterization of NPM1 condensates from CG simulations. **a)** Representative bright field and mCherry fluorescence images of single condensate at increasing protein concentrations (1–20 µM). Condensate size visibly increases with higher mCherry-NPM1 levels. **b)** Scatter plot of surface area (bright field) versus fluorescence intensity (mCherry) for condensates at 1 µM and 20 µM, illustrating a clear shift toward larger and brighter particles at higher concentration. **c)** Quantification of surface area from 10,000 individual condensates across protein concentrations, showing a concentration-dependent increase in average condensate size. **d)** Snapshot of NPM1 slab configuration (100 chains) from CG simulation in which two phases coexist. Red, green and blue colored segments represent the N-terminal OD domain, the IDR region and the C-terminal domain respectively of the full-length NPM1 protein (*Top*). Density profile of NPM1 along the z-dimension of the slab geometry. The area between grey dashed lines denotes the dense phase and the area outside of it represents the dilute phase (*Bottom*). **e)** Time-averaged intermolecular contact map of NPM1 proteins within the condensed phase. The contact number of NPM1 is normalized to the highest contact number of NPM1. Each colored bar highlights the corresponding domain of the NPM1 protein. The panel below the 2D contact map shows the average number of contacts per chain as a function of residue number. Statistical analysis was performed using unpaired, two-tailed student’s *t*-tests. Significance is denoted as follows: p < 0.0001 (****). Scale bars = 7 µm.

### Molecular dynamics studies of NPM1

While these experimental approaches provide crucial insight into the morphology of NPM1 condensates, they do not directly reveal how intramolecular architecture contributes to phase separation. Specifically, it remains unclear how domain-specific interactions within NPM1 drive condensate formation and what role individual NPM1 domains play in self-association. Previous studies have largely focused on how NPM1 interacts with RNA or other macromolecular partners, leaving gaps in our understanding of how NPM1 alone can organize into phase-separated assemblies.^45–47^ To address this, we performed coarse-grained (CG) molecular dynamics simulations to investigate how structural domains within NPM1 contribute to self-association and condensate formation.

NPM1 is a modular protein composed of three domains: an *N*-terminal oligomerization domain (OD, residues 1–118, red), a central intrinsically disordered region (IDR, residues 119–242, green), and a *C*-terminal DNA-binding domain (CTD, residues 243–294, blue) (**Fig. S3a-b**). AlphaFold-based structure predictions revealed high-confidence folding (pLDDT > 90) for the OD and CTD, and low pLDDT scores for the IDR, indicating a flexible and disordered central region (**Fig. S3c**). Notably, charge analysis revealed striking asymmetries: the OD and IDR are highly negatively charged, with net charges of –7 and –19 respectively, while the CTD is mildly positively charged (+2), consistent with its known affinity for nucleic acids (**Fig. S4a**). Despite their shared negative charge, these domains differ in charge patterning. Using the Sequence Charge Decoration (SCD) parameter, which quantifies local charge clustering in disordered regions,^48^ we found that the IDR region has the most charge segregation (SCD = –1.473), with negative and positive charges clustered within the IDR segment (**Fig. S4a-b**). Such clustering can enhance phase separation,^49, 50^ thus motivating further exploration of how these features can potentially govern self-assembly of NPM1.

To understand the role of these domain-specific properties in NPM1 phase separation, we simulated 100 full-length NPM1 chains using the HPS-Urry model^51^ under slab geometry^52, 53^ to achieve coexisting dense and dilute phases. We applied rigid body motion to folded domains (OD and CTD), while disordered regions remained fully flexible (see Supporting Information for simulation details). We simulated this system at a fixed temperature of 300 K with 100 mM salt and analyzed protein densities as a function of the z-coordinate (**Fig. 4d**). The density profile clearly shows that the system phase separates into distinct dense and dilute phases, with NPM1 condensates forming a well-defined dense phase (**Fig. 4d**). To probe the molecular interactions that mediate condensate formation, we computed 2D and 1D time-averaged intermolecular contact maps across all domains (**Fig. 4e**).

Despite both the OD and IDR being negatively charged, the OD region exhibited the strongest intermolecular contacts, particularly OD–OD interactions (**Fig. 4e** and **Fig. S4c**). Many residues in this region formed the highest number of intermolecular contacts (see lower panel in **Fig. 4e**). This aligns with its structural role in forming stable pentamers, as seen in the crystal structure of human NPM1 (PDB: 4N8M), and suggests that it acts as a scaffold for multivalent network formation. Interestingly, the IDR also displayed much stronger IDR–IDR interactions, despite its strong negative charge (see **Fig. S4c**). This observation highlights the importance of charge patterning over net charge, as the presence of segregated blocks of charges within the IDR may facilitate transient attractions and enable phase separation. In contrast, CTD–CTD interactions were weak (**Fig. S3c**), consistent with its function as a nucleic acid-binding region rather than a driver of protein–protein condensation. However, we observed favorable CTD–IDR and CTD–OD interactions, suggesting that the CTD may help stabilize the condensate by bridging domains or enhancing multivalency.

Together, these findings demonstrate that *in vitro* condensate formation can be reliably assessed using both traditional methods and newer approaches such as flow cytometry. Notably, imaging flow cytometry (IFC) confirms that mCherry-NPM1 condensates retain their droplet-like morphology under flow, validating the structural integrity of particles analyzed in this format. In parallel, our CG simulations provide complementary molecular-level insight into the organization of NPM1 within the condensed phase. The domain-specific interactions, not directly accessible via experimental imaging alone, highlight the molecular architecture underpinning condensate stability. By integrating high-throughput flow-based methods with molecular modeling, we establish a multidimensional platform for quantifying condensate size, morphology, and internal organization, providing a powerful framework for dissecting the physical basis of biomolecular phase separation.

### Colocalization study of NPM1 condensates with macromolecules

Having established the formation and characterization of NPM1 condensates, we next sought to investigate their ability to recruit and interact with various macromolecules. Biomolecular condensates often serve as hubs for molecular organization, selectively incorporating proteins, nucleic acids, and small molecules based on physicochemical properties such as charge, hydrophobicity and multivalency.^23, 54–57^ To assess the partitioning behavior of different macromolecules within NPM1 condensates, we systematically examined the interaction of NPM1 condensates with antibodies, lipids, small-molecule drugs, and RNA (**Fig. 5**).

**Fig. 5.**
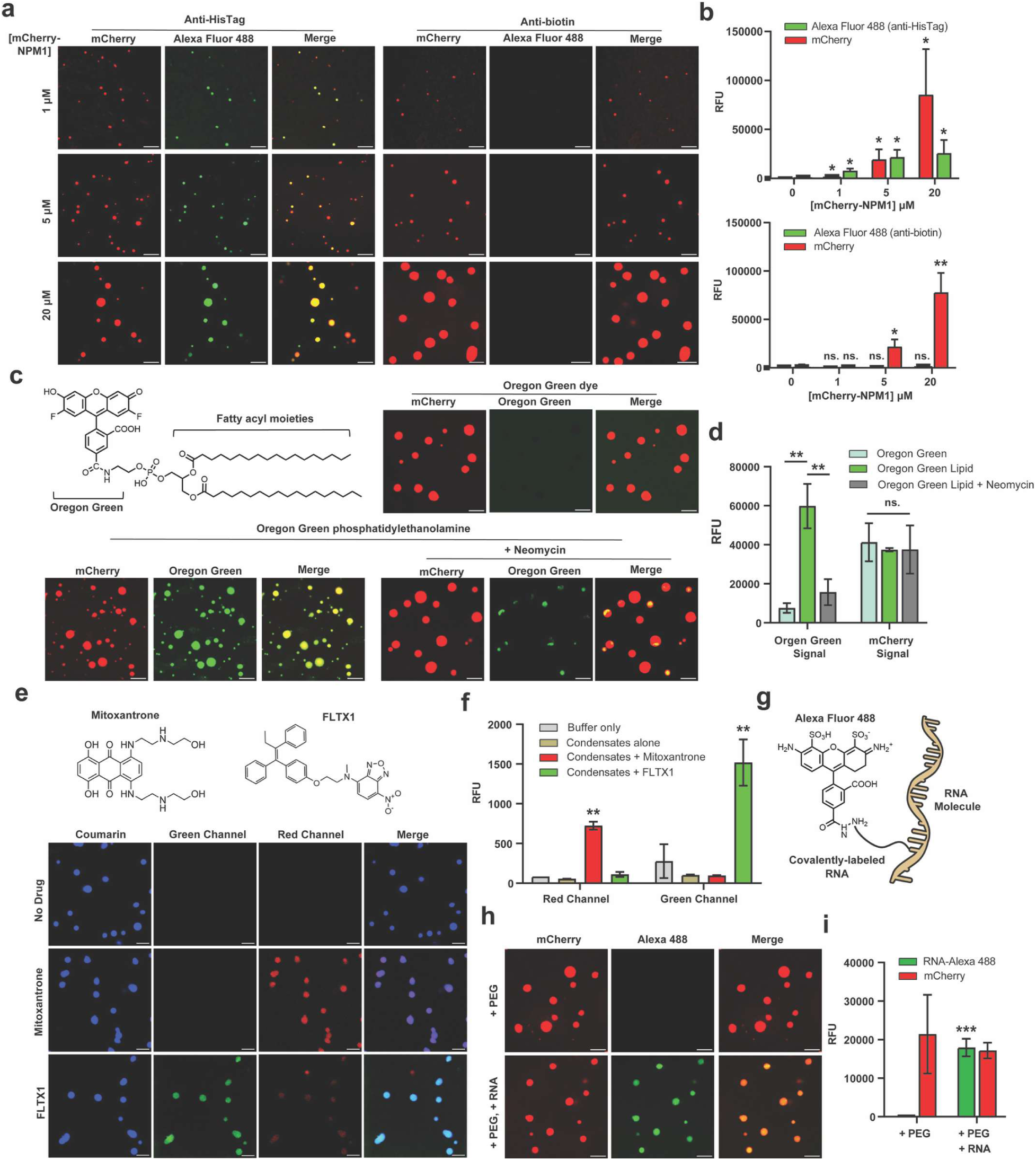
Colocalization study of NPM1 condensates with macromolecules. **a)** Confocal imaging of mCherry-NPM1 condensates incubated with Alexa Fluor 488-labeled anti-HisTag (positive control) and anti-Biotin (negative control) antibodies at different NPM1 concentrations. Anti-HisTag is selectively recruited into NPM1 condensates, while anti-biotin remains excluded. **b)** Flow cytometry confirms colocalization results from **a)**, showing increased Alexa Fluor 488 fluorescence for anti-HisTag but not for anti-Biotin. Statistical comparisons are made against the 0 µM protein condition **c)** Confocal imaging of 20 µM NPM1 condensates incubated with 2 µM lipid probes reveals selective partitioning of Oregon Green phosphatidylethanolamine (a lipid-modified dye) but exclusion of Oregon Green 488 (a hydrophilic dye), indicating a preference for lipid-like molecules. **d)** Flow cytometry corroborates the imaging data in **c)**, showing increased Alexa Fluor 488 fluorescence in the presence of Oregon Green phosphatidylethanolamine. Pre-incubation with 2 mg/mL neomycin disrupts lipid association without affecting mCherry-NPM1 condensate formation. **e)** Confocal imaging of 20 μM NPM1 condensates incubated with small-molecule drugs (50 μM mitoxantrone or 100 μM FLTX1) demonstrates their strong partitioning into condensates. **f)** Flow cytometry confirms drug partitioning, showing an increase in fluorescence intensity for both mitoxantrone and FLTX1 in NPM1 condensates. Statistical comparisons are between mitoxantrone versus buffer only (red channel), and FLTX1 versus buffer only (green channel). **g)** Schematic representation of RNA labeling using Alexa Fluor 488. **h)** Confocal imaging shows strong colocalization of RNA-Alexa 488 with NPM1 condensates, indicating RNA recruitment. **i)** Flow cytometry analysis supports confocal imaging in **h)**, demonstrating robust RNA partitioning into NPM1 condensates. Statistical comparisons are between condensates with RNA and condensates without RNA. Statistical analysis was performed using unpaired, two-tailed student’s *t*-tests. Error bars indicate standard deviation (SD). Significance is denoted as follows: p < 0.05 (*), p < 0.01 (**), p < 0.001 (***), p < 0.0001 (****) and not significant (ns.) for p≥ 0.05. Scale bars = 5 µm.

To start, we performed confocal imaging of NPM1, which contains a terminal His-tag fusion, using Alexa Fluor 488-labeled anti-HisTag (positive control) and anti-biotin (negative control) antibodies. Higher NPM1 concentrations led to larger condensates in both the mCherry and Alexa Fluor 488 channels (**Fig. 5a**), indicating successful recruitment of the anti-HisTag antibody. Flow cytometry mirrored these colocalization profiles, showing a clear increase in both mCherry and Alexa Fluor 488 intensity (**Fig. 5b**). In contrast, the anti-biotin antibody did not partition into NPM1 condensates, as evidenced by the absence of Alexa Fluor 488 signal in both confocal imaging and flow cytometry (**Fig. 5a-b**). Together, these results demonstrate selective antibody partitioning and establish flow cytometry as a complementary, quantitative approach for characterizing condensate–macromolecule interactions.

Recent studies have demonstrated that phospholipids can partition into condensates, potentially influencing their size, morphology, and function.^58^ In our study, we performed colocalization assays using two lipid probing dyes: Oregon Green 488 (a hydrophilic dye) and Oregon Green phosphatidylethanolamine (a lipid-conjugated dye with a hydrophobic tail). Confocal imaging revealed that Oregon Green 488 did not colocalize with NPM1 condensates, as only mCherry fluorescence was observed, suggesting that the condensates exclude freely soluble small molecules (**Fig. 5c**). In contrast, Oregon Green phosphatidylethanolamine strongly partitioned into NPM1 condensates, indicating that lipid-modified molecules can associate with these structures. These results mirror the previously described behavior of these two molecules with biomolecular condensates using confocal microscopy.^58^ We paired these experiments with flow cytometry analysis and, once again, found that flow cytometry can yield results that mirror the general trend but that have a much greater sample pool size (**Fig. 5d**). In these experiment, Alexa 488 fluorescence was low when NPM1 was incubated with Oregon Green but significantly increased when incubated with Oregon Green phosphatidylethanolamine, demonstrating its strong partitioning into NPM1 condensates. However, when NPM1 was pre-incubated with neomycin before adding Oregon Green phosphatidylethanolamine, the Alexa 488 signal remained low, suggesting that neomycin disrupts lipid association, likely by interfering with electrostatic or hydrophobic interactions.^59^ Meanwhile, mCherry fluorescence intensity remained high across all conditions (**Fig. 5d**), indicating that condensate formation itself was not affected. These findings support the idea that biomolecular condensates can selectively incorporate lipid-like molecules, potentially modulating their biochemical properties.

Likewise, prior research also highlighted that small-molecule drugs can selectively partition into biomolecular condensates, potentially influencing their local concentration and pharmacodynamics.^21^ To benchmark this feature in a high-volume analysis platform, we performed colocalization assays using mitoxantrone and FLTX1, two small-molecule drugs with distinct chemical properties.^60, 61^ Confocal imaging confirmed that both mitoxantrone and FLTX1 strongly partitioned into NPM1 condensates, as evidenced by their respective intrinsic fluorescence signals in the red and green channels (**Fig. 5e**).

These findings were further supported by flow cytometry analysis, which showed a clear increase in drug-associated fluorescence within condensates (**Fig. 5f**). These results suggest that NPM1 condensates can serve as selective compartments for small-molecule therapeutics, highlighting the potential impact of condensate partitioning on drug distribution and activity.

Finally, we investigated the association of RNA with NPM1 condensates, given the well-documented role of RNA in nucleating and modulating condensate formation in various cellular contexts.^9^ We labeled total RNA extracted from *E. coli* with AZDye 488 (**Fig. 5g**) and incubated it with mCherry-NPM1 in the presence of PEG8000. Confocal imaging showed strong colocalization of mCherry and AZDye 488 signals, indicating that RNA colocalizes with NPM1 condensates (**Fig. 5h**). Flow cytometry analysis further confirmed the localization of the two biomacromolecules (**Fig. 5i**), demonstrating robust RNA association with NPM1-driven condensates. We also incubated RNA with mCherry-NPM1 in the absence of PEG. While condensates were still formed (**Fig. S5**), they were noticeably smaller and less spherical compared to those formed with PEG (**Fig. 5h**). This result suggests that while RNA alone can promote NPM1 condensate formation, the resulting structures are substantially smaller, highlighting the role of crowding agents like PEG in enhancing condensate growth and detectability *in vitro*. Collectively, these colocalization studies demonstrate that NPM1 condensates selectively recruit a wide range of macromolecules, highlighting their potential as compartmentalized microenvironments for molecular interactions. The complementary use of flow cytometry not only validated imaging observations but also revealed subtle shifts in condensate composition, underscoring its utility as a reliable and scalable approach for dissecting the molecular content of biomolecular condensates.

### Assessment of biomolecular condensate dynamics

Having benchmarked flow cytometry across a range of established contexts, we next aimed to uncover features of biomolecular condensates that require both finer temporal resolution and high-throughput data acquisition. To this end, we focused on the dynamic behavior of individual proteins and their potential partitioning between the dilute and dense phases. Biomolecular condensates are highly dynamic structures^62^ that continuously exchange molecules with their surroundings, yet quantitatively capturing these dynamic events remains a major challenge in the field. Traditional approaches, such as confocal microscopy and fluorescence recovery after photobleaching (FRAP), have provided valuable insights into molecular mobility within condensates,^46, 63, 64^ but they are often limited in throughput and time resolution, making it difficult to assess real-time exchange events between condensates.

To investigate exchange dynamics between condensates, we replaced mCherry with HaloTag to generate an NPM1–HaloTag fusion protein. During affinity purification, we installed three distinct chloroalkane-linked fluorescent dyes – Coumarin, R110, or TAMRA – enabling site-specific labeling of the fusion protein. While the protein was bound to the resin, fluorescent dyes were conjugated, and excess unreacted dye was subsequently washed away. This strategy ensured full removal of the unbound dye (**Fig. S6**). We next performed *in vitro* droplet formation assays using either individually prepared dye-labeled NPM1 variants or mixtures of differentially labeled variants and monitored condensate morphology over time via confocal microscopy (**Fig. 6a**). At time zero, condensates displayed distinct, non-overlapping fluorescence signals. After 30 minutes, however, most droplets exhibited complete fluorescence colocalization, indicative of molecular exchange and droplet fusion (**Fig. 6a**). While these results clearly demonstrate rapid molecular exchange between condensates composed of the same protein, capturing this dynamic process in real time remains challenging with confocal microscopy due to its limited temporal resolution.

**Fig. 6.**
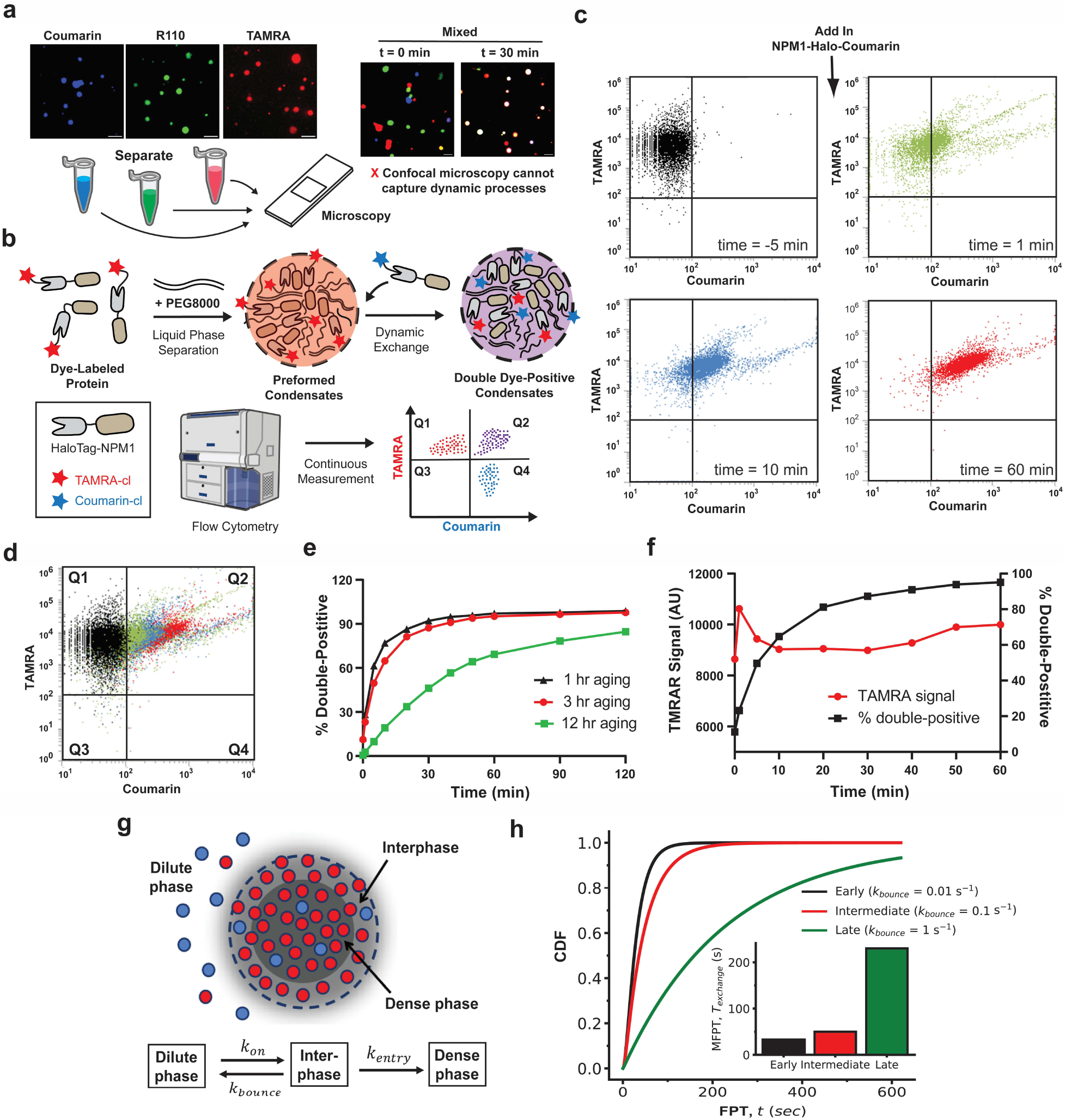
Assessment of biomolecular condensate dynamics. **a)** Confocal imaging of 10 µM NPM1-Halo condensates labeled with distinct fluorescent dyes (Coumarin, R110, or TAMRA) shows that at t = 0, droplets remain separate, but after 30 minutes, they merge, indicating molecular exchange. However, confocal imaging lacks the temporal resolution to capture this rapid process. Scale bars = 5 µm. **b)** Schematic of the flow cytometry-based assay for continuous real-time measurement of condensate exchange in a single-tube format, unlike conventional 96-well plate methods. Pre-formed NPM1-Halo-TAMRA (red) condensates were mixed with NPM1-Halo-Coumarin (blue) protein solution, and fluorescence exchange was monitored over time. **c)** Flow cytometry scatter plots tracking molecular exchange. Initially, preformed NPM1-Halo-TAMRA condensates exhibit a homogeneous scatter in the TAMRA^+^/Coumarin^-^ quadrant. Over time, events gradually shift toward the double-positive (TAMRA^+^/Coumarin^+^) window, indicating progressive molecular exchange. **d)** Overlay of all scatter plots from panel **c** shows a continuous and directional population shift toward the double-positive quadrant. **e)** Quantification of exchange dynamics for 1-hour, 3-hour, and 12-hour aged condensates. Younger condensates rapidly incorporate new proteins (>97% double-positive events within 2 hours), while older condensates show slower exchange, with 12-hour aged condensates reaching only ∼80%, suggesting increased rigidity or reduced mobility with aging. **f)** Monitoring of TAMRA fluorescence in 3-hour aged condensates reveals a slight decrease after blue protein addition, likely due to transient redistribution or structural rearrangements. **g)** Schematic of the discrete-state stochastic model for blue protein exchange into a preformed red condensate. The model considers three spatially distinct discrete states: the dilute phase (surrounding solution), the interfacial layer (condensate surface), and the dense phase (droplet interior). Proteins transition from dilute to interphase with rate 𝑘_on_, return to dilute phase with rate 𝑘_bounce_, or proceed into the dense phase with rate 𝑘_entry_ . **h)** Theoretical cumulative distribution functions (CDFs) of exchange times computed under different aging conditions by varying 𝑘_bounce_. Inset: Mean first-passage time (MFPT) of exchange for early (𝑘_bounce_ = 0.01 𝑠^-1^), intermediate (𝑘_bounce_ = 0.1 𝑠^-1^) and late or aged (𝑘_bounce_ = 1.0 𝑠^-1^) condensates. Other parameters used for the calculations are: 𝑘_on_ = 0.05 𝑠^-1^ and 𝑘_entry_ = 0.1 𝑠^-1^.

To further confirm that fluorescence accumulation within condensates reflects specific molecular interactions rather than nonspecific partitioning of small molecules, we performed a control using small fluorescent HaloTag ligands. Pre-formed NPM1-Halo condensates were incubated with either TAMRA-chloroalkane, which covalently binds HaloTag, or a structurally similar TAMRA-alkane lacking HaloTag-binding capability. Flow cytometry analysis revealed strong TAMRA fluorescence within condensates only in the presence of TAMRA-chloroalkane, whereas TAMRA-alkane produced no detectable signal (**Fig. S7**). These results demonstrate that small molecules without specific binding affinity do not spontaneously accumulate within NPM1-Halo condensates, supporting the conclusion that observed fluorescence redistribution in our system is driven by defined molecular interactions.

We leveraged flow cytometry to quantitatively analyze exchange dynamics between condensates at the single-droplet level with high sensitivity and throughput. The ability of flow cytometry to rapidly measure fluorescence intensity across thousands of droplets in solution enables real-time tracking of molecular transfer between condensates. To accomplish this, we designed an assay (**Fig. 6b**) in which NPM1-Halo-TAMRA (red) condensates were pre-formed by equilibration with PEG8000. Subsequently, we introduced an NPM1-Halo-Coumarin (blue) protein solution lacking PEG and immediately monitored fluorescence over time by flow cytometry. We designated -5 min as the time point corresponding to pre-formed red condensates before mixing, and 0 min as the point of blue protein addition.

To confirm that the observed increase in blue fluorescence within red condensates was due to active molecular exchange rather than passive mixing or dye diffusion in the bulk solution, we performed a control using free TAMRA dye. A TAMRA-only solution (without protein) was added to PEG8000 at the same volume ratio used in the exchange assay. After a brief 2-second mixing period, the dye rapidly dispersed and became uniformly distributed throughout the PEG solution (**Fig. S8**), confirming that unbound fluorophores equilibrate freely in the solution phase. This result supports the conclusion that the observed colocalization in our primary assay arises specifically from incorporation of NPM1-Halo-Coumarin into pre-existing condensates, rather than from nonspecific mixing of fluorescent components.

We observed that introducing NPM1-Halo-Coumarin into pre-formed NPM1-Halo-TAMRA condensates resulted in rapid molecular exchange, with over 40% of events shifting to the double-positive quadrant (TAMRA⁺/Coumarin⁺) immediately at time zero. This rapid exchange raised concerns that critical transient events might be missed due to the high rate of molecular transfer. To investigate whether condensate dynamics could be artificially slowed, we explored the effect of condensate aging. Prior studies have shown that NPM1 condensates readily undergo fusion within 60 minutes of incubation but exhibit markedly reduced fusion rates after 180 minutes, likely due to aging-associated structural changes.^65^ Motivated by these observations, we pre-incubated NPM1-Halo-TAMRA condensates with PEG8000 for 1, 3, or 12 hours at room temperature prior to the addition of NPM1-Halo-Coumarin. We then monitored 10,000 individual events over time using quadrant-based flow cytometry analysis to assess changes in exchange dynamics.

As expected, at -5 minutes (prior to the addition of NPM1-Halo-Coumarin) pre-formed NPM1-Halo-TAMRA condensates exhibited a homogeneous scatter distribution within the TAMRA⁺/Coumarin⁻ quadrant (**Fig. 6c**). Upon addition of NPM1-Halo-Coumarin, we observed a progressive shift of events toward the double-positive TAMRA⁺/Coumarin⁺ quadrant, indicating active molecular exchange between the two protein pools (**Fig. 6c-d**). Raw scatter plots corresponding to these distributions are presented in **Fig. S9**. Quantification of the percentage of double-positive events over time revealed that the exchange rate declined with increasing aging time of the pre-formed condensates (**Fig. 6e**), suggesting that condensate dynamics diminish as they mature. To assess how the introduction of NPM1-Halo-Coumarin impacts the structural integrity of aged NPM1-Halo-TAMRA condensates, we monitored the TAMRA fluorescence intensity in 3-hour aged samples. Within 60 minutes of blue protein addition, the TAMRA signal exhibited a slight but consistent decrease (**Fig. 6f**). This reduction likely reflects the redistribution of red-labeled NPM1 due to molecular exchange. Initially, the red condensates were in equilibrium with their environment; however, the sudden influx of fresh blue protein may have perturbed this balance, resulting in transient dilution or internal rearrangements that lowered fluorescence intensity.

To further support this interpretation, we performed a control experiment in which preformed NPM1-Halo-TAMRA condensates were mixed with an equal concentration of soluble NPM1-Halo-TAMRA protein lacking PEG (**Fig. S10**). Notably, even in this NPM1-Halo-TAMRA followed by NPM1-Halo-TAMRA condition, we observed a similar slight decrease in overall TAMRA fluorescence within 60 minutes. This suggests that the observed drop in fluorescence is not specific to dye identity or heterotypic interactions but may arise from protein influx-induced redistribution or structural remodeling of the condensates.

For the 1-hour and 3-hour aged samples, molecular exchange was highly efficient, with approximately 97% of condensates becoming double-positive within 2 hours of mixing (**Fig. 6e**). In contrast, the 12-hour aged sample reached only ∼80% double-positive, indicating that condensate aging markedly reduces exchange efficiency. This diminished capacity may result from increased structural rigidity, reduced molecular mobility, or fibrillation-like transitions that impede protein incorporation. Supporting this, confocal imaging of 3-hour aged condensates after 2 hours of mixing with the blue protein showed predominantly spherical morphologies. In contrast, 12-hour aged condensates exhibited irregular, broken shapes (**Fig. S11**), consistent with aging-induced structural changes that impair condensate dynamics.

To further gain mechanistic insights into the dynamics of protein exchange during aging, we developed a discrete-state stochastic model that describes transitions of individual blue proteins across three spatially distinct states: the dilute phase (surrounding solution), the interfacial layer, and the dense phase (droplet interior) of a preformed red condensate, as shown schematically in **Fig. 6g**. This interfacial region plays a crucial role in controlling exchange kinetics, because it acts as a kinetic barrier for molecular entry.^66^ In our theoretical model, blue proteins initially located in the dilute phase can enter the interfacial region with rate 𝑘_on_, return or bounce back to the dilute phase with rate 𝑘_bounce_ , or proceed inward from interphase into the dense phase of the red condensate with rate 𝑘_entry_ (see **Fig. 6g**). To describe the exchange dynamics of blue proteins into a preformed red condensate, we used a method of first-passage probabilities. More specifically, one can define a function 𝐹_dil_(𝑡) as the probability density of a blue protein entering the dense phase of the red condensate for the first time at time 𝑡 if initially at 𝑡 = 0, the system started in the dilute phase. Similar first-passage probability density function for the interphase can be defined as 𝐹_int_(𝑡). The temporal evolution of these probabilities is governed by the backward master equations^67^:

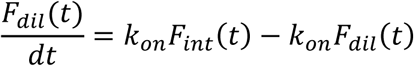

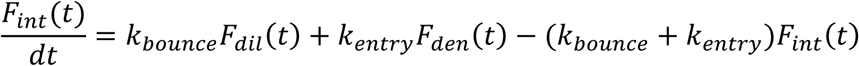

where 𝐹_den_(𝑡) is the probability density of a blue protein being found in the dense phase immediately after leaving the interphase. So, it is natural to assume that 𝐹_den_(𝑡) = 𝛿(𝑡). This physically means that if the system is in this state at 𝑡 = 0, the process is immediately finished. As described in detail in the supporting information, these equations can be solved analytically for all ranges of parameters using Laplace transformations 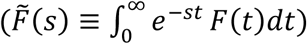 , producing the first-passage time (FPT) distribution 𝐹(𝑡) for entering the dense phase,

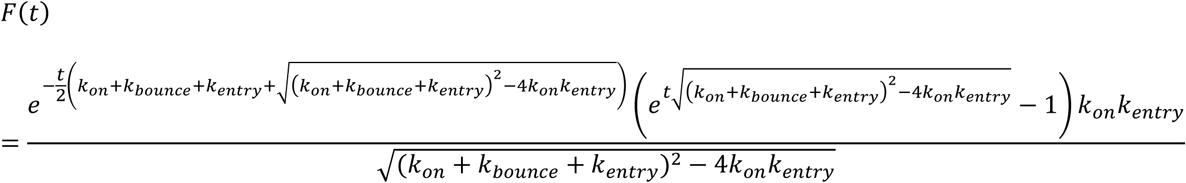

We also obtained the corresponding mean first-passage time (MFPT) for protein exchange, as given by

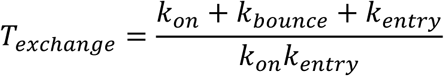

Because the experimental setup introduces blue proteins into solution while red droplets are preformed, the system is inherently nonequilibrium as droplets can grow by recruiting newly arrived proteins. We therefore adopted kinetic rate constants from a nonequilibrium theoretical framework described by Wurtz and Lee,^68^ where chemical reaction-controlled exchange governs droplet formation and dynamics.

We then examined how aging affects molecular exchange by modulating 𝑘_bounce_, which controls how readily proteins leave the interfacial region before entering the dense phase.

This parameter is particularly sensitive to interfacial structural properties. In a recent experiment by Emmanouilidis *et al.* ^69^ that combined NMR and Raman spectroscopies with microscopy, droplet maturation or aging were investigated for FUS protein, where the structural properties upon maturation between the inside and the surface of droplets were obtained. It was found that a solid crust-like shell is observed at the surface that comprised of 𝛽-sheet contents and ultimately matured droplets were converted into fibril-like structures. This suggests that due to aging or maturation of condensates, the interface of the droplet can be much more coarsened due to this structural appearance and therefore exchange upon aging can be limited by the dynamics at the droplet interface, implying the existence of an interface resistance.^66, 70^ Based on this observations of interfacial solidification, we considered three conditions: “early” (low interfacial resistance, 𝑘_bounce_ = 0.01 𝑠^-1^ ), “intermediate” ( 𝑘_bounce_ = 0.1 𝑠^-1^ ), and “late or matured” condensates (high resistance, 𝑘_bounce_ = 1.0 𝑠^-1^ ). As shown in the inset of **Fig. 6h**, exchange becomes progressively slower with aging, with MFPT increasing from ∼32 𝑠 (early) to ∼230 𝑠 (late), in agreement with experimental delays in double-positive accumulation (**Fig. 6e**). Moreover, additional structural insights from Chatterjee *et al.*^71^ show that interfacial regions become kinetically trapped in 𝛽-sheet–like conformations, further slowing dynamics compared to the untrapped interior. We mimicked this by decreasing 𝑘_entry_ in our model for aged droplets, which also led to slower exchange times (see **Fig. S12**). To further support this connection, we computed the cumulative distribution function (CDF) from the first-passage distribution:

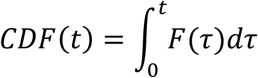

which represents the probability that a protein has entered the dense phase by time 𝑡, and thus serves as a proxy for the fraction of blue proteins colocalized within red condensates. Under different aging conditions, the CDF curves (**Fig. 6h** and **Fig. S12**) show a clear trend: rapid rise for early, young condensates and gradual rise for late, aged ones—qualitatively recapitulating the experimental curves in **Fig. 6e**. Despite the differing timescales (theoretical FPT is in seconds and experimental mixing time is in minutes), the alignment between the theoretical and experimental dynamics strongly supports that younger condensates support faster exchange, while aged condensates accumulate proteins more slowly due to interfacial resistance and decreased internal mobility.

To test whether condensate dynamics are temperature-sensitive, we performed the protein exchange assay using pre-formed, red-labeled condensates (3 h aged) and kept on ice. Upon addition of the blue-labeled protein, minimal incorporation was observed over 90 minutes, with a low percentage of double-positive droplets detected by flow cytometry (**Fig. S13**). However, when the same sample was returned to room temperature, a rapid increase in double-positive events was observed, eventually reaching levels comparable to our standard room temperature assay. These results suggest that lower temperatures suppress dynamic exchange, likely by reducing molecular mobility within aged condensates.

To further demonstrate the versatility of our condensate dynamics assay, we extended its application beyond protein exchange to monitor the incorporation of macromolecular cargo. Specifically, we tested whether lipid molecules could be dynamically recruited into pre-formed condensates by mixing NPM1-Halo-TAMRA condensates with Oregon Green-conjugated phosphatidylethanolamine (OG-PE). Flow cytometry analysis revealed a progressive increase in the double-positive population over 30 minutes (**Figure S14**), indicating successful incorporation of the fluorescent lipid into the red condensates. This observation was supported by the migration of events toward the TAMRA(+)/OG(+) quadrant in the scatter plots (**Figure S14**). These results highlight that our real-time flow cytometry platform can sensitively track condensate dynamics involving diverse molecular components, offering a broadly applicable tool to uncover new principles of phase separation and inform future biological investigations.

## CONCLUSION

In this study, we established a multi-modal framework to investigate the structure, composition, and dynamics of biomolecular condensates. Using NPM1 as a model scaffold protein, we demonstrated that flow cytometry can serve as a quantitative, high-throughput method for analyzing condensate formation and macromolecular partitioning, complementing traditional imaging-based approaches. Imaging flow cytometry (IFC) confirmed the structural integrity of condensates under flow, validating the reliability of flow-based analysis.

To further probe the molecular basis of NPM1 condensate formation, we performed coarse-grained molecular dynamics simulations that uncovered the role of domain-specific interactions in driving NPM1 self-association. Our simulations revealed that the N-terminal oligomerization domain (OD) of NPM1 engages in strong homotypic interactions, while the intrinsically disordered region (IDR), despite its net negative charge and charge-segregated sequence, also contributes extensively to condensate stability. In contrast, the C-terminal DNA-binding domain (CTD) exhibited weaker self-interactions but formed favorable contacts with both OD and IDR regions. These findings highlight how architectural features and sequence-level electrostatics coordinate the self-assembly of NPM1 condensates in the absence of nucleic acids or other binding partners.

We established a flow-based molecular exchange assay, which revealed that condensate aging reduces exchange efficiency, likely due to altered biophysical properties such as internal organization and reduced mobility. To provide mechanistic insight into these experimental findings, we developed a discrete-state stochastic model that describes exchange as a sequence of transitions between dilute, interfacial, and dense regions of a condensate. This framework captures the kinetics of single-protein exchange events using first-passage time distributions and mean exchange times. By varying key kinetic parameters - specifically the rate of escape from the interface and the entry rate into the dense phase - we recapitulated the experimentally observed slowing of exchange with condensate aging. Our theoretical model further predicts cumulative exchange behavior *via* first-passage time analysis, which closely aligns with the experimental time courses of double-positive droplet accumulation (indicating successful molecular exchange between condensates) across aging conditions. Together, these theoretical results provide a quantitative framework that complements and explains our experimental observations, highlighting the role of interfacial resistance and molecular mobility in aging-dependent molecular exchange dynamics.

Compared to traditional microscopy-based approaches, our integrated workflow enables more quantitative and mechanistic insight into condensate behavior. This platform can be extended to investigate other condensate systems, including how crowding agents, chaperones, or small molecules influence condensate dynamics and stability. While flow cytometry offers robust quantification and scalability, it also presents certain challenges that inform these future applications. Condensates are dynamic and sensitive to environmental changes, and shear stress or buffer composition within flow systems may subtly influence their behavior or stability. Moreover, distinguishing between true condensates and nonspecific aggregates remains difficult without additional validation steps, such as imaging or enzymatic assays. By integrating experimental assays with molecular simulations and theoretical modeling, these limitations can be effectively mitigated. This comprehensive approach enables deeper and more reliable investigation into the dynamic and structural properties of biomolecular condensates.

## ACKNOWLEDGEMENT

This study was supported by the NIH grants 1R01AI178975-01 (M.M.P.), R35GM124893 (M.M.P.), R35GM153388 (J.M.), R01AI179080-01 (M.M.P.). We thank the W.M. Keck Center for Cellular Imaging for the usage of Zeiss LSM 980 microscopy System and Leica STELLARIS 8 confocal/FLIM/tauSTED microscope system (NIH OD030409) and thank the Flow Cytometry Core Facility for the usage of Amnis ImageStreamX Mark II system. We acknowledge the Texas A&M High Performance Research Computing (HPRC) for providing computational resources that have contributed to the results reported in the article. The content of this work is solely the responsibility of the authors and does not necessarily represent the official views of the NIH.

## SUPPORTING INFORMATION

Additional figures, tables, and materials/methods are included in the supporting information file.

