## Supplementary Information for "A High-Throughput, Flow Cytometry Approach to Measure Phase Behavior and Exchange in Biomolecular Condensates"

**Fig. S1**

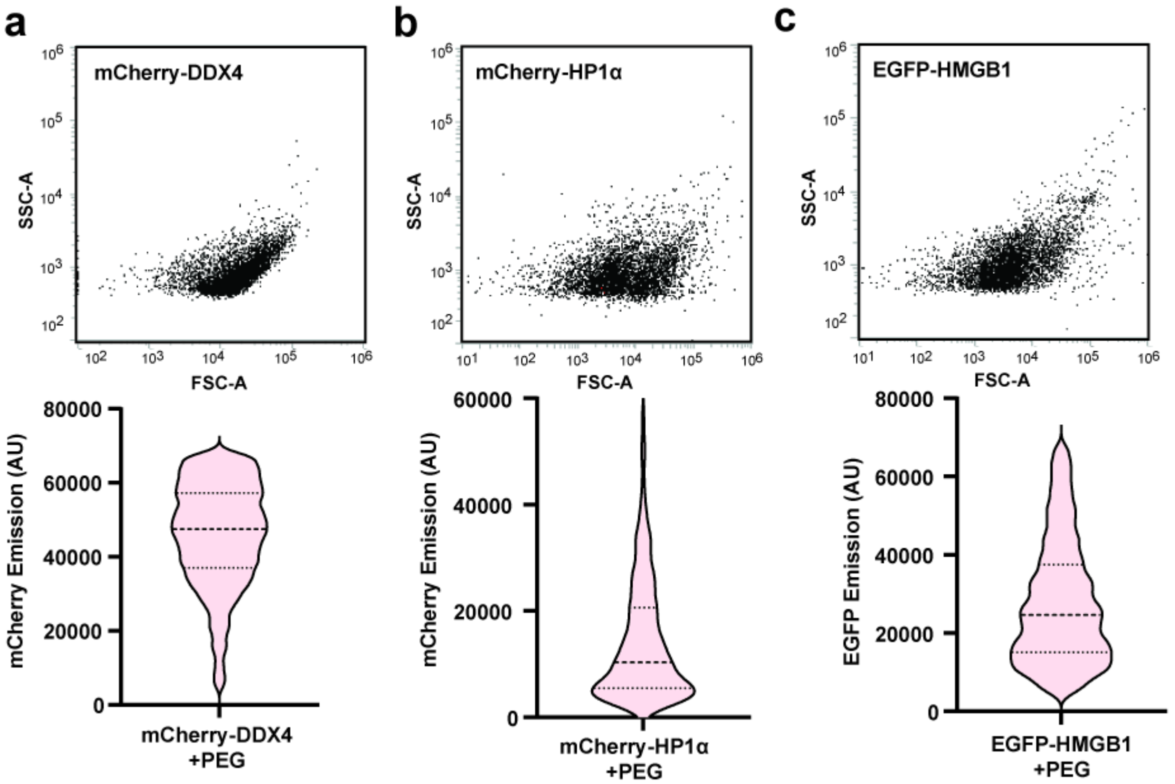

**Flow cytometry characterization of PEG-induced condensates for additional scaffolding proteins. a)** top: Side scatter area (SSC-A) versus forward scatter area (FSC-A) plot for 20 µM mCherry-DDX4 in the presence of 10% PEG8000, reflecting condensate size and granularity. *Bottom:* Violin plot of fluorescence intensity distribution for 5,000 gated condensates. **b)** Same analysis as in **a)**, performed for 20 µM mCherry-HP1α. **c)** Same analysis as in **a)**, performed for 20 µM EGFP-HMGB1. These results demonstrate that flow cytometry robustly captures both particle complexity and fluorescence heterogeneity across different protein condensates.

**Fig. S2**

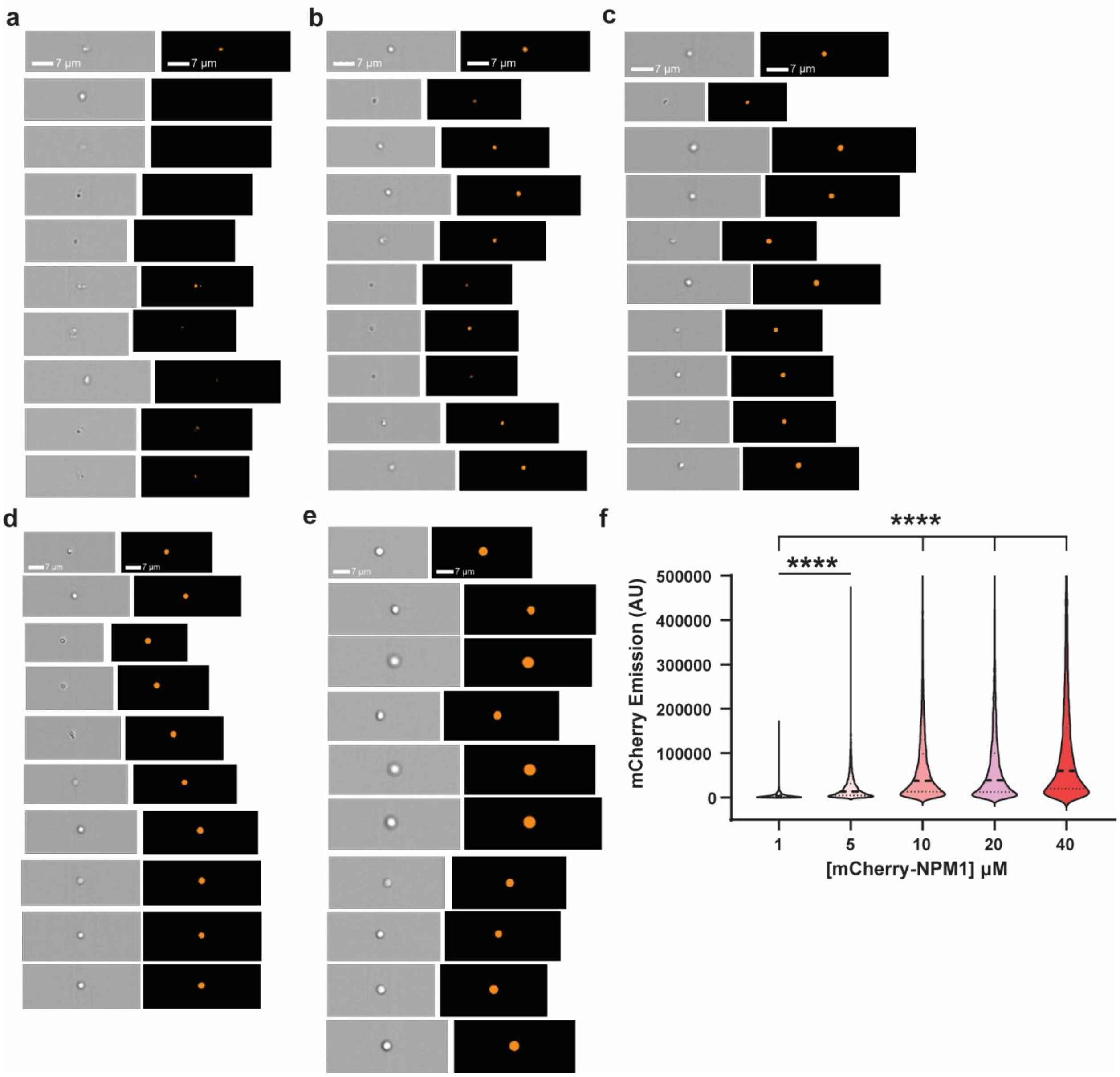

**Additional representative images and quantification of mCherry-NPM1 condensates captured by imaging flow cytometry (IFC). a-e)** Representative IFC images of mCherry-NPM1 condensates formed at increasing concentrations (1, 5, 10, 20, and 40 µM) in the presence of 10% PEG. Each panel includes images from both the bright field and mCherry fluorescence channels. A general increase in condensate size is observed with higher protein concentrations. Scale bar = 7 µm. **f)** Violin plot showing the distribution of mCherry fluorescence intensity for over 10,000 condensates at each protein concentration. The fluorescence intensity increases with protein concentration, consistent with increased condensate formation. Statistical analysis was performed using unpaired, two-tailed student's *t*-tests. Significance is denoted as follows: $p < 0.0001$ (****).

Fig. S3

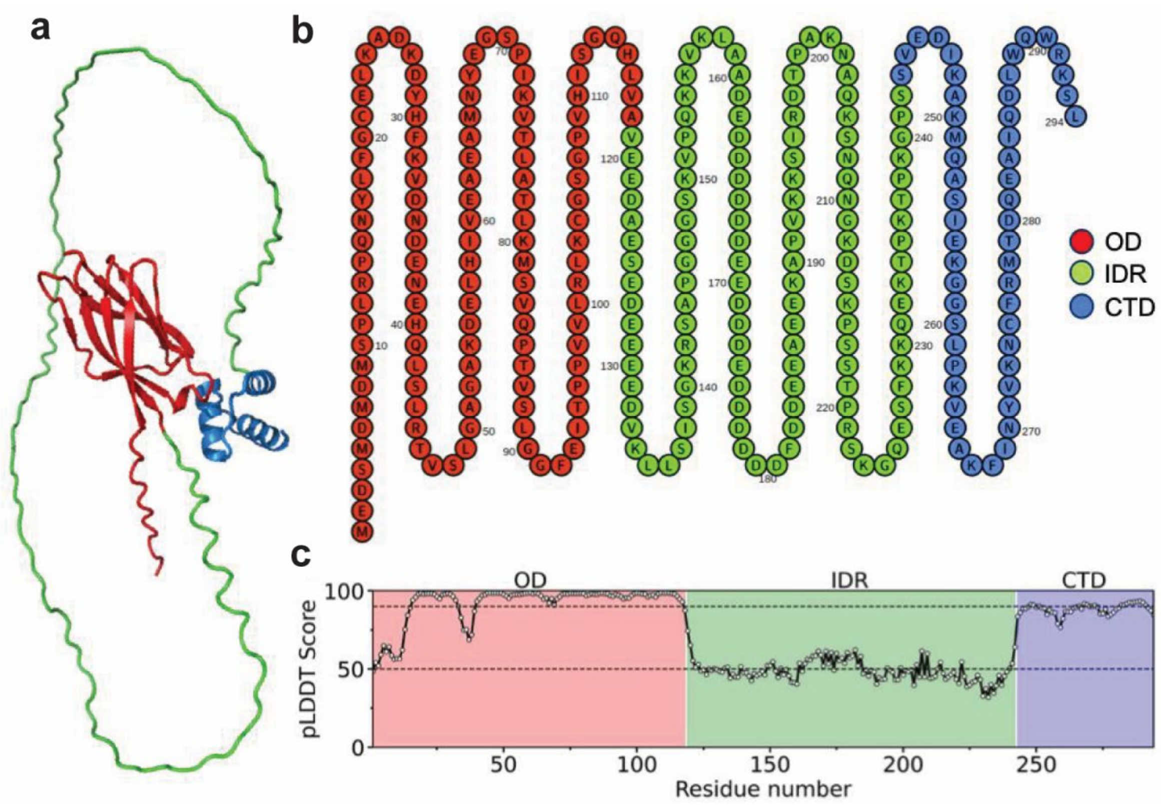

**Structure and sequence of NPM1 protein used in coarse-grained (CG) simulation. a)** Cartoon representation of the Alpha-Fold generated structure NPM1 protein. The N-terminal oligomerization domain (OD) and the C-terminal domain (CTD) of NPM1 are shown by red and blue colors respectively, while the central intrinsically disordered region (IDR) is shown by green color. **b)** Sequence of the full-length NPM1 protein. **c)** pLDDT (predicted local distance difference test) scores for each residue of the AlphaFold model of NPM1 protein. High pLDDT score (above 90, indicated by black dotted line) indicates high confidence in the prediction, in which both the backbone and side chains are typically predicted with high accuracy, where low pLDDT score (below 50, indicated by black dotted line) indicates a region that is naturally highly flexible or intrinsically disordered.

Fig. S4

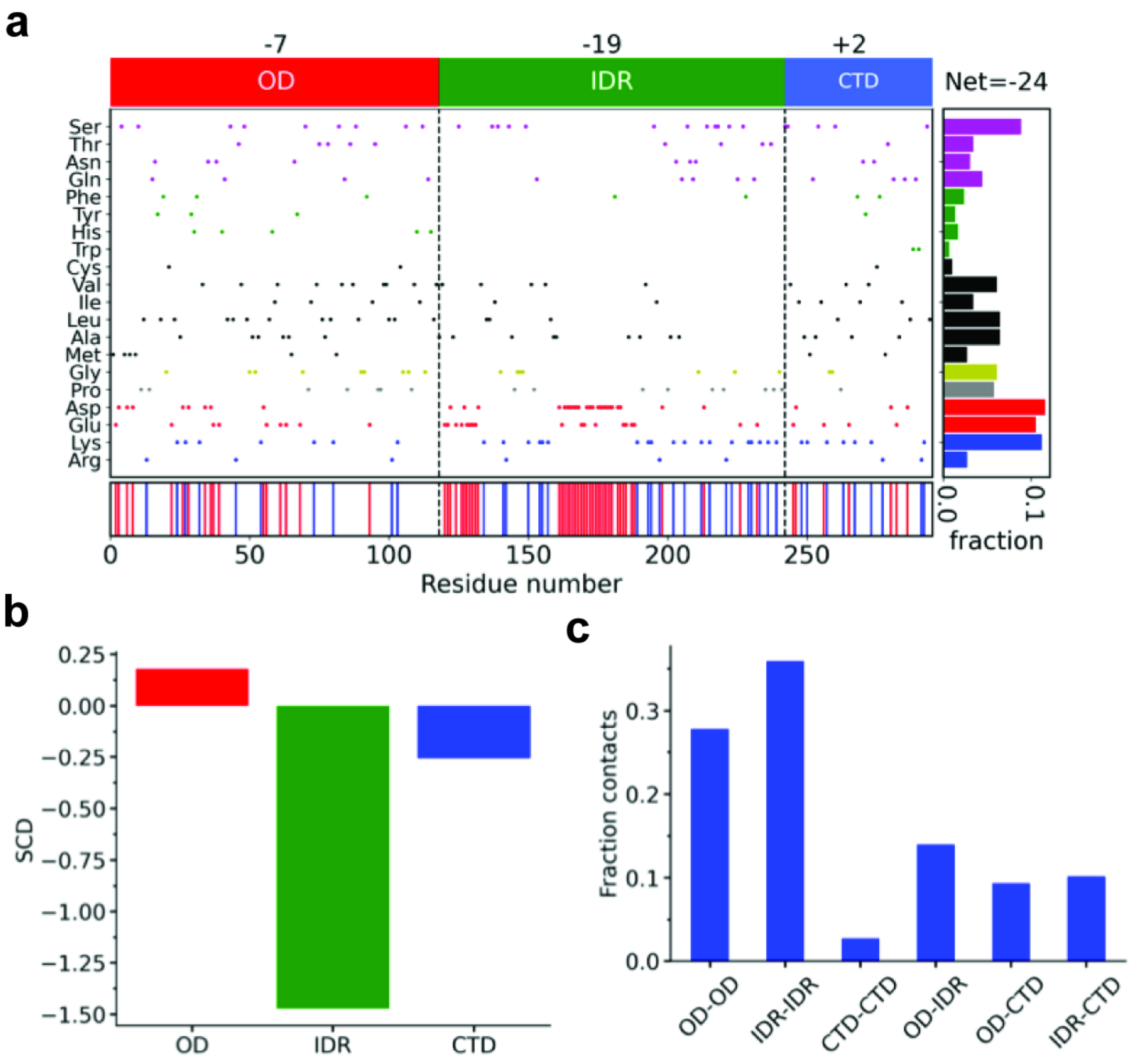

**Sequence and domain-level charge organization and contact propensity in NPM1 condensates. a)** Domain architecture of the full length NPM1 protein, showing the net charge of each domain and distribution of amino acids. Red and blue vertical lines indicate the location of negatively and positively charged residues. **b)** SCD (sequence charge decoration) values for the OD, IDR and CTD regions of NPM1 protein, highlighting the degree of charge segregation.[1] A large negative value of SCD indicates a block-like charge arrangement (which is reflected in the IDR domain in the above panel), while a value close to 0 reflects an alternating charge pattern along the sequence. **c)** Fractional contributions of domain-domain intermolecular contacts (OD–OD, IDR–IDR, CTD–CTD, OD–IDR, OD–CTD, and IDR–CTD) obtained from coarse-grained co-existence simulations of NPM1 condensates.

Fig. S5

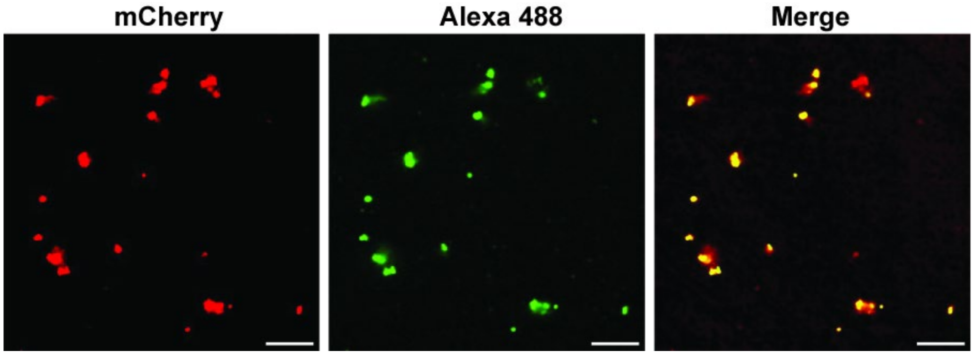

**RNAs alone can induce condensate formation**. Confocal images of mCherry-NPM1 (20 μM) incubated with RNA (50 ng/μL) in the absence of PEG. In this condition, mCherry-NPM1 forms visible condensate-like assemblies, though their morphology appears irregular and less spherical compared to condensates formed under crowding conditions.

**Fig. S6**

**a** **b** **c** **d**

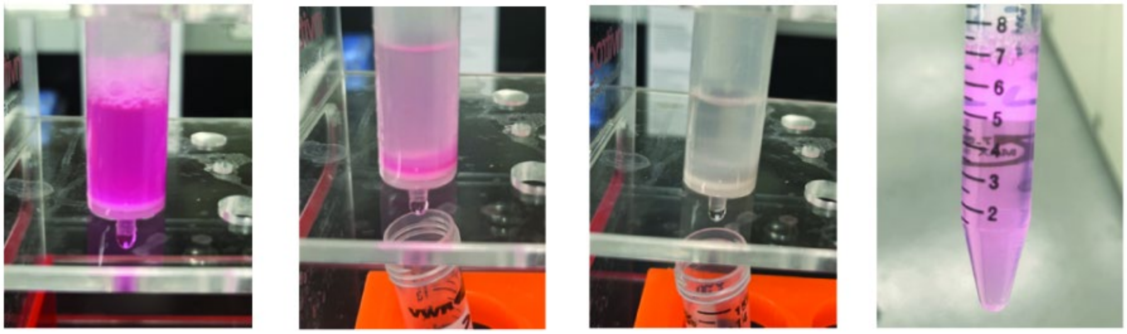

**Photographic documentation of the purification process of TAMRA-labeled NPM1-Halo proteins. a)** During the binding step, NPM1-Halo proteins covalently labeled with TAMRA-chloroalkane (TAMRA-Cl) exhibit a distinct red color on the Ni-NTA resin, indicating successful dye conjugation. **b)** As the wash step proceeds under gravity flow, the red color fades, reflecting the removal of excess unbound dye. **c)** Upon elution with buffer containing high imidazole, most labeled proteins are released from the resin, and the resin becomes nearly colorless. d) The final eluate containing TAMRA-labeled NPM1-Halo proteins appears visibly red in the collection tube, confirming successful labeling and purification. NPM1-Halo proteins labeled with R110-chlooalkane or Coumarin-chloroalkane were purified using the same protocol.

**Fig. S7**

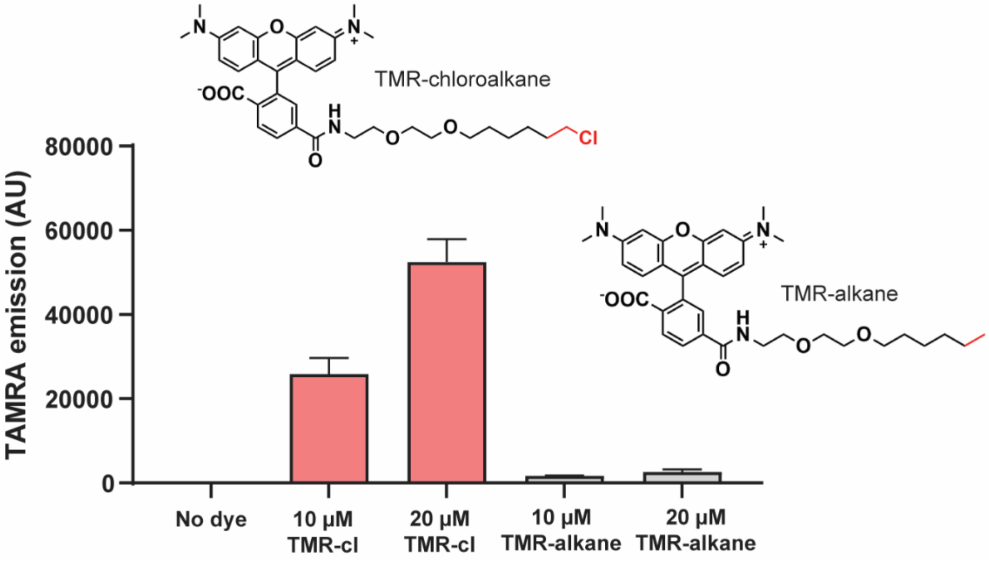

**TAMRA-chloroalkane but not TAMRA-alkane accumulates in NPM1-Halo condensates.** NPM1-Halo protein (20 µM) was incubated with 10% PEG8000 for 30 min to form condensates, then treated with either TAMRA-chloroalkane (TMR-Cl) or TAMRA-alkane (TMR-alkane) for 30 min. Flow cytometry showed strong, concentration-dependent fluorescence with TMR-Cl, but minimal signal with TMR-alkane, indicating selective accumulation through specific HaloTag binding.

**Fig. S8**

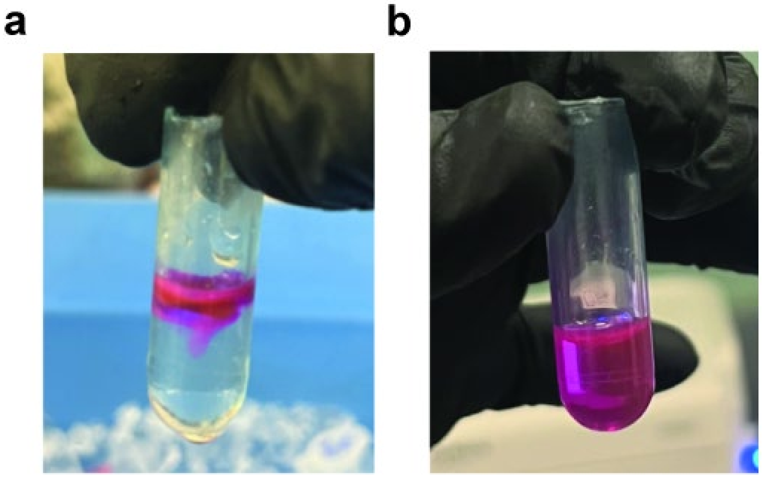

**Assessment of TAMRA dye mixing in PEG-containing solutions. a)** TAMRA-chloroalkane (TAMRA-Cl) dye was added to a 10% PEG solution that had been incubated at room temperature for 3 hours. Without agitation, the dye remained largely undissolved and floated on top of the solution even after 10 minutes, indicating poor spontaneous mixing. **b)** After a brief 2-second vortexing at maximum speed, the dye became fully dispersed, showing homogeneous mixing throughout the PEG solution. These observations confirm that in our condensate dynamic exchange assays, the signal detected by flow cytometry reflects genuine molecular exchange between pre-formed condensates and labeled proteins, rather than incomplete mixing of proteins or dyes in solutions.

**Fig. S9**

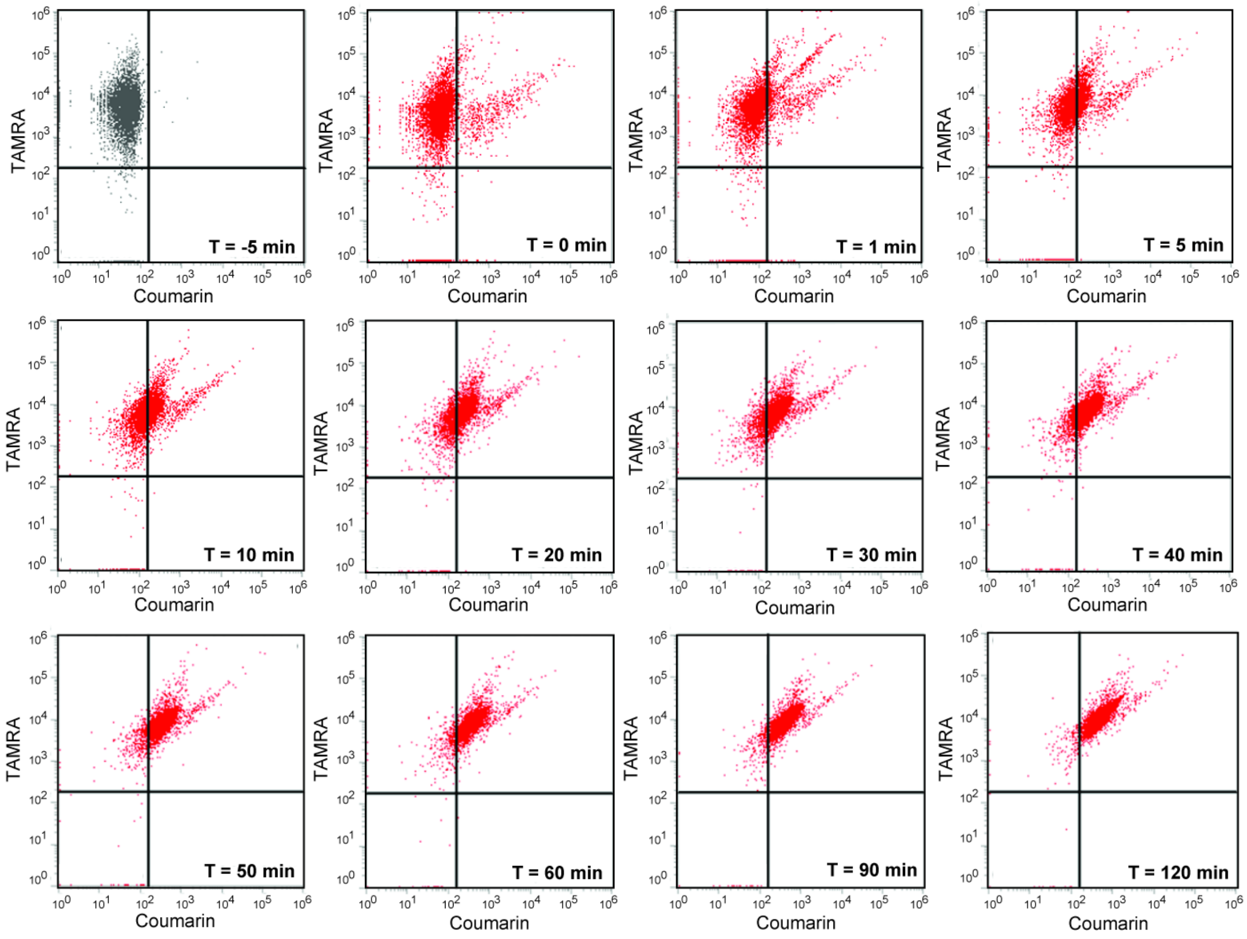

**Flow cytometry scatter plots show dynamic exchange between pre-formed NPM1-Halo-TAMRA condensates and incoming NPM1-Halo-Coumarin protein.** 10 μM pre-formed NPM1-Halo-TAMRA condensates (3 hr aged) were mixed with 10 μM soluble NPM1-Halo-Courmain protein (no PEG) at time 0 min. At T = -5 min, events appear mostly in the TAMRA-positive quadrant, indicating homogeneous red condensates. Upon mixing, populations gradually shift toward the double-positive quadrant (TAMRA+/Coumarin+), reflecting protein exchange over time. By 120 min, over 95% of events reside in the double-positive quadrant, demonstrating efficient exchange which can be captured by flow cytometry.

**Fig. S10**

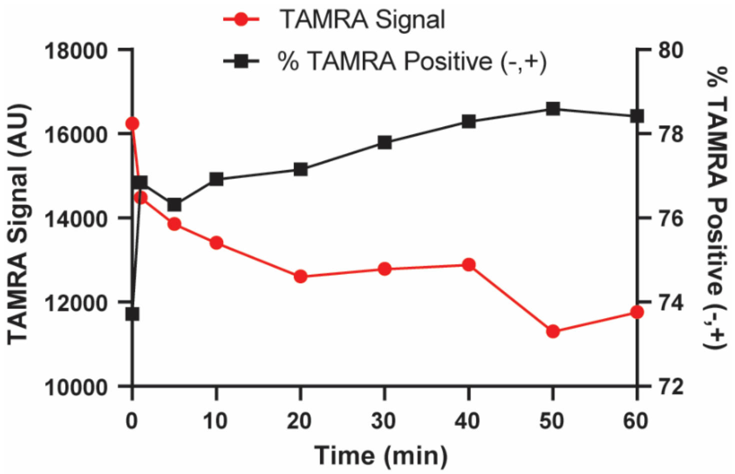

**Flow cytometry detects dynamic exchange between pre-formed NPM1-Halo-TAMRA condensates and soluble NPM1-Halo-TAMRA protein.** 10 μM pre-formed NPM1-Halo-TAMARA condensates (3 hr aged) were mixed with 10 μM soluble NPM1-Halo-TAMRA protein and analyzed over time. Due to identical fluorophores, only the TAMRA-positive (-,+) quadrant was tracked. A slight increase in event count suggests incorporation of soluble protein into condensates. The modest drop in mean TAMRA fluorescence may result from fluorophore dilution, photobleaching, or environmental quenching upon exchange.

**Fig. S11**

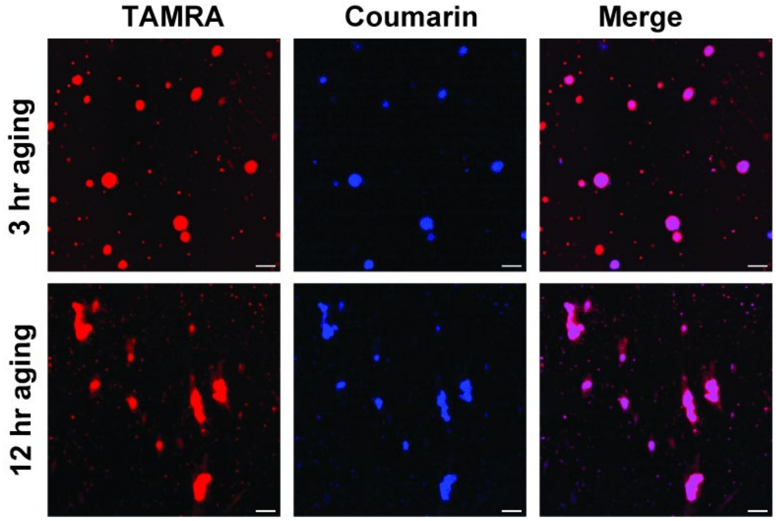

**Morphological differences in NPM1-Halo condensates following dynamic exchange assays at different aging times.** Confocal images show the structural integrity of NPM1-Halo-TAMRA condensates before and after dynamic exchange with NPM1-Halo-Coumarin. **Top row**: 3-hour aged NPM1-Halo-TAMRA condensates maintain a generally round morphology and structural integrity following exchange with NPM1-Halo-Coumarin, indicating a more dynamic and fluid-like state. **Bottom row**: 12-hour aged NPM1-Halo-TAMRA condensates show disrupted morphology and the presence of aggregate-like structures after the same exchange treatment, suggesting increased rigidity or solid-like properties with longer aging.

**Fig. S12**

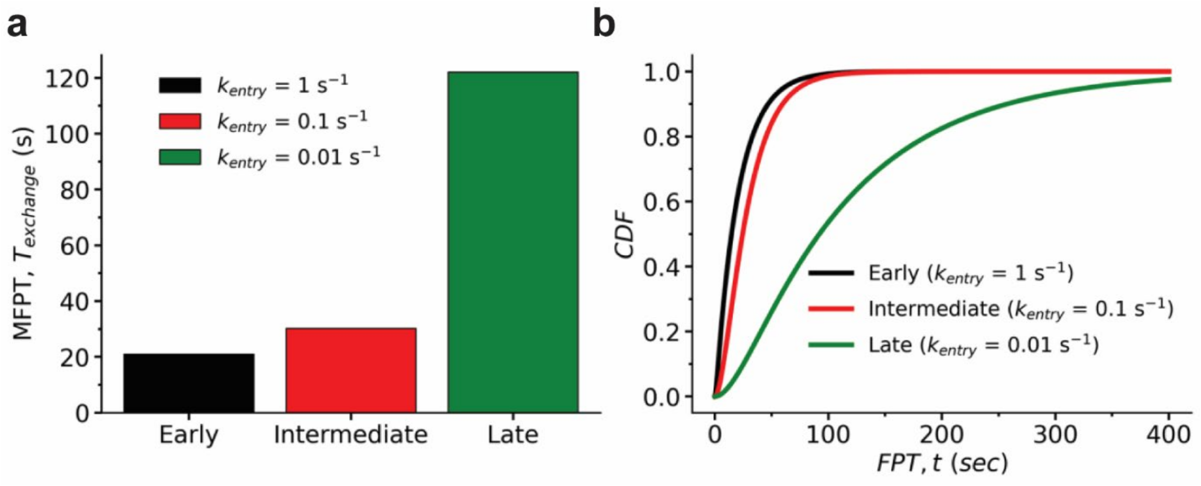

**Impact of droplet aging via $k_{entry}$ on protein exchange dynamics. a)** Mean first-passage time (MFPT) of exchange for early ($k_{entry}$=1 $s^{-1}$), intermediate ($k_{entry}$=0.1 $s^{-1}$) and late or aged ($k_{entry}$=0.01 $s^{-1}$) condensates. **b)** Theoretical cumulative distribution functions (CDFs) of exchange times computed under different aging conditions by varying $k_{entry}$. Other parameters used for the calculations are: $k_{on}$=0.05 $s^{-1}$ and $k_{bounce}$=0.001 $s^{-1}$.

**Fig. S13**

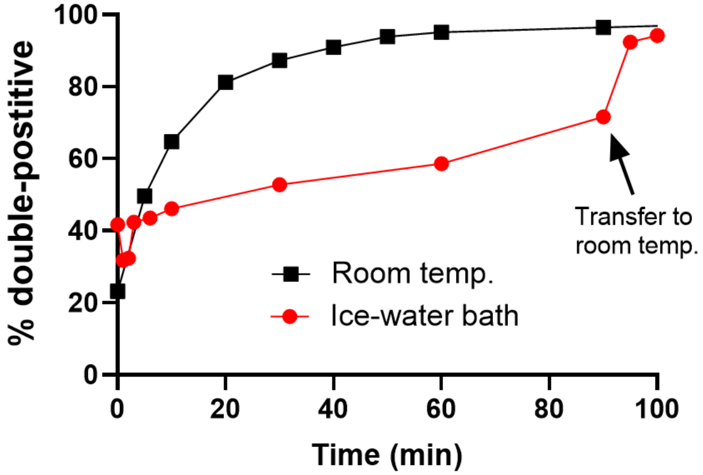

**Temperature-dependent exchange dynamics within aged NPM1 condensates.** Pre-formed NPM1-Halo-TAMRA condensates (10 μM, 3 hr aged) were mixed with 10 μM soluble NPM1-Halo-Coumarin protein and monitored over time using flow cytometry. Samples kept on ice showed a slow increase in double-positive droplets (%), as indicated by the red curve. After 90 minutes, returning the sample to room temperature led to a rapid increase in the double-positive population. For comparison, the black curve represents the same exchange assay performed for a 3 hr aged sample at room temperature (data from Figure 6e). Both conditions eventually reached >90% double-positive droplets, indicating temperature-dependent modulation of exchange dynamics and eventual equilibration.

**Fig. S14**

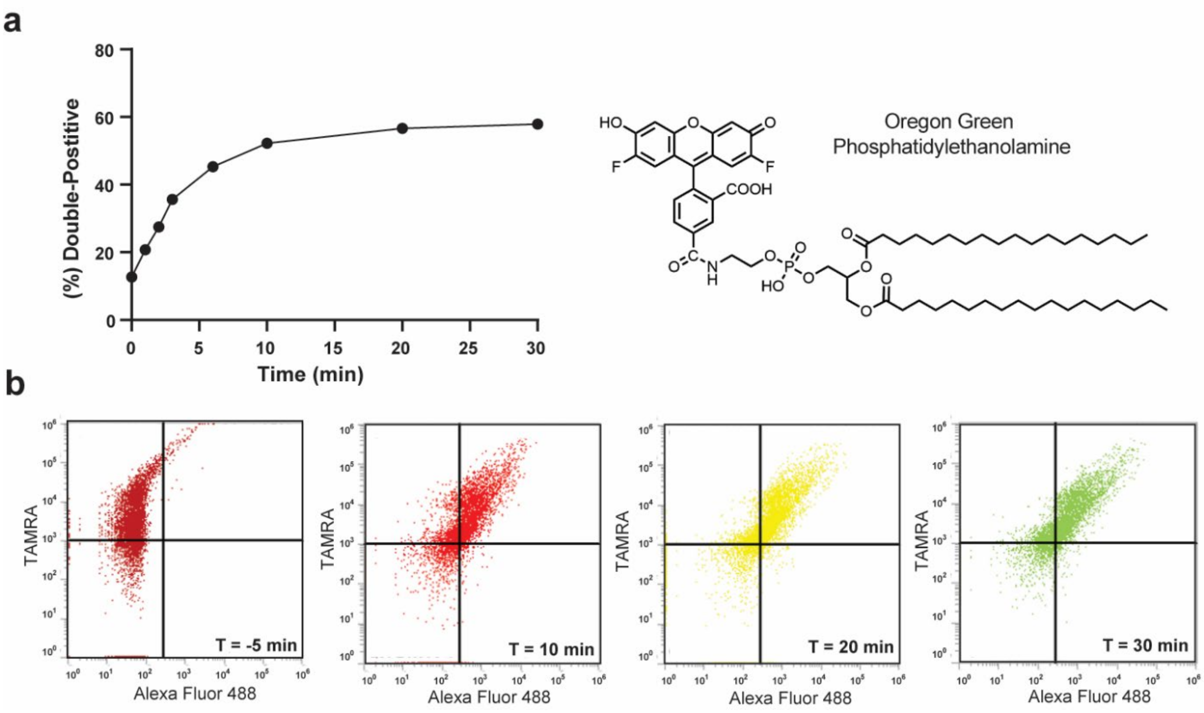

**Flow cytometry enables real-time tracking of lipid molecule incorporation into pre-formed condensates. a)** Quantification of double-positive populations over time during dynamic exchange between pre-formed NPM1-Halo-TAMRA condensates and Oregon Green-conjugated phosphatidylethanolamine (OG-PE). b) Representative scatter plots showing the dynamic progression of molecular exchange. A clear migration of condensate populations into the TAMRA(+)/Alexa Fluor 488(+) quadrant is observed over 30 minutes after OG-PG addition.

### General Experimental Details

*Instrumentation and Reagents*

DNA concentrations were quantified using a NanoDrop spectrophotometer (Thermo Fisher, CHEM-PR1-KIT). Images from agarose gels and SDS-PAGE were captured with a ChemiDoc XRS+ gel imager (Bio-Rad, 1708265). A Synergy$^{TM}$ H1 hybrid multi-mode reader (Agilent, BioTek Synergy H1) was employed for condensate turbidity measurement. Confocal images were obtained on a Zeiss LSM 980 Microscopy System and a Leica STELLARIS 8 confocal/FLIM/tauSTED microscope system equipped with tunable white light laser. For most flow cytometry analyses related to condensates, an Attune$^{TM}$ NxT Acoustic Focusing Cytometer was used. For imaging flow cytometry (IFC) analysis, the Amnis ImageStreamX Mark II system (Luminex Corporation) was employed.

Protein expression was carried out in *Escherichia coli* BL21-Gold (DE3) competent cells (Agilent Technologies, 230132). RNA extraction was performed using *Escherichia coli* BL21 (DE3) pLysS cells. A Monarch Total RNA Purification Kit (New England Biolabs, T2010S) was used to facilitate RNA extraction from bacterial cultures. Protein expression and purification involved the use of Isopropyl-β-D-thiogalactopyranoside (IPTG) (Chem Impex, 00194), cOmplete protease inhibitor cocktail (Millipore Sigma, 11836153001), Econo-Pac Columns (Bio-Rad, 7321010), HisPur$^{TM}$ Ni-NTA resin (Thermo Fisher, 88221), Amicon Ultra-15 Centrifugal Filter Unit (EMD Millipore, UFC900324), Pierce™ Slide-A-Lyzer® Dialysis Cassettes (Thermo Fisher, 66380). Chemicals used for protein and RNA labeling, as well as macromolecule compounds for condensate colocalization studies, were obtained from commercial sources. AZDye$^{TM}$ 488 hydrazide (Vector Laboratories, FP-1017) was used to label bacterial RNA. Polyethylene Glycol 8000 (Sigma-Aldrich, PHR2894-1G) was used to induce condensate formation, and 1,6-Hexanediol (Sigma-Aldrich, 240117-50G) was used to disrupt condensates. Flow cytometry assays were conducted in 96-well untreated round-bottom plates (VWR, 82050-622).

*Protein Expression and Purification*

The plasmids encoding scaffolding proteins were acquired from Addgene: pET-45b-mCherry-NPM1 (Addgene, 194546), pET28C-mCherry-DDX4LCD (Addgene, 204408), pET-45b-mEGFP-HMGB1-WT (Addgene, 194543), pET-45b-HP1a-mCherry (Addgene, 185012). The above plasmids were a gift from Denes Hnisz from Max Planck Institute, and Samie Jaffrey from Cornell

University. Codon-optimized DNA sequences of HaloTag fusion scaffolding protein (NPM1-Halo) were synthesized by GenScript and cloned into a pET-28a vector with an N-terminus His-tag. All plasmids were transformed into *Escherichia coli* BL21-Gold (DE3) cells. The transformed cells were then inoculated into Luria-Bertani (LB) media and shaken at 37°C until $OD_{600}$ reached 0.3-0.6. Protein expression was induced by adding 1 mM isopropyl β-D-1-thiogalactopyranoside (IPTG), followed by incubation at 37°C for 3 hr with shaking at 250 rpm. This step was followed by an extended incubation at 18°C for 18 hr. Cells were harvested by centrifugation at 4000 rpm for 30 min. The resulting cell pellets were lysed via sonication in lysis buffer (50 mM Tris-Cl, pH 7.5 at 25°C; 1 M NaCl; 1 mM DTT) supplemented with protease inhibitors (Sigma-Aldrich, 11836153001). The lysates were centrifuged at 18,000 g for 20 min at 4°C, and the clarified supernatant was applied to Ni-NTA resins (Thermo Fisher, 88221) in a gravity flow column (Bio-Rad, 7321010). The resin was allowed to bind the proteins at 4°C overnight with rotation. The column was washed with 100 mL of wash buffer (50 mM Tris-Cl, pH 7.5 at 25°C; 1 M NaCl; 1 mM DTT; 20 mM imidazole). Recombinant proteins were eluted with 10 mL elution buffer (50 mM Tris-Cl, pH 7.5 at 25°C; 500 mM NaCl; 500 mM imidazole). The eluted proteins were concentrated using an ultra-15 centrifugal filter unit (EMD Millipore, UFC901024), and dialyzed into 1X storage buffer (50 mM Tris-Cl, pH 7.5 at 25°C; 125 mM NaCl; 1 mM DTT; 10% glycerol). Protein concentrations were determined using a NanoDrop spectrophotometer (Thermo Fisher, 13-400-518), based on the calculated extinction coefficient and molecular weight of each protein. Corresponding protein sequences are provided in **Table S1**.

*Protein Labeling*

Our lab has previously established protocols to synthesize chloroalkane ligand dyes for HaloTag binding.[2] Nonetheless, for pursuing more consistent results, we chose to purchase dyes from commercially available vendors: TAMRA-Chloroalkane (TAMRA-Cl, Promega, G8252), Coumarin-Chloroalkane (Coumarin-Cl, Promega, G8582), and R110-Chloroalkane (R110-Cl, Promega, G3221). These dyes were directly conjugated to the NPM1-Halo protein while it was bound to Ni-NTA resin, enabling efficient on-resin labeling. Briefly, clarified lysate containing NPM1-Halo was loaded onto 1 mL of Ni-NTA resin in a gravity flow column. After washing with 20 mL of wash buffer to remove non-specific proteins, the resin was incubated for 15 minutes at room temperature with 5 mL of 1X storage buffer containing 30 µL of 10 mM stock dye (TAMRA-Cl, Coumarin-Cl, or R110-Cl) under gentle shaking. Following labeling, the column was washed with 100 mL of wash buffer to remove unbound dye, and labeled proteins were eluted using 10 mL of elution buffer. Eluted proteins were then concentrated and dialyzed following the same protocol described in the purification section above.

*RNA Isolation and Labeling*

*Escherichia coli* BL21 (DE3) pLysS cells were grown at 37°C in LB media until $OD_{600}$ reached 1.0, then centrifuged at 4000 g for 30 min. All subsequent steps were performed at room temperature to prevent RNA precipitation. RNA isolation was carried out using the Total RNA Extraction kit (NEB, T2010) following the manufacturer's protocol. In brief, cells were resuspended in 1X DNA/RNA Protection Reagent with 1 mg/mL lysozyme (MP Biomedicals, 100831), followed by sonication for 30 min. The lysates were transferred to an RNase-free microfuge tube and centrifuged at 16,000 g for 2 min. The supernatant containing RNA was mixed with RNA Lysis Buffer and processed according to the manufacturer's instructions. The eluted RNA was further concentrated using isopropanol and 70% ethanol, and the final RNA concentration was determined using NanoDrop. For RNA labeling, we were using an established similar protocol from previous reference.[3] Briefly, 87 µL of RNA (3.0 mg/mL) was mixed with 3.3 µL of 3 M sodium acetate (pH 5.2) and 10 µL of freshly prepared 25 mM sodium periodate. The mixture was incubated on ice for 50 min, followed by the addition of 20 µL of 3 M sodium acetate (pH 5.2) and 80 µL of nuclease-free water. RNA was precipitated by adding 400 µL of isopropanol and incubating on ice for 1 h. The RNA was pelleted by centrifugation at 16,000 g for 15 min at 4°C, then rinsed with 150 µL of ice-cold ethanol. After another centrifugation at 16,000 g for 15 min, the RNA pellets were resuspended in 500 µL of reaction buffer containing 100 mM sodium acetate (pH 5.2) and 25 nmol AZDye 488 Hydrazide (Vector Laboratories, FP-1017-1). The reaction was incubated for 48 hr at room temperature with gentle shaking. The labeled RNA was purified via additional isopropanol and ethanol precipitation and dissolved in 100 µL of nuclease-free water. The final RNA-AZDye488 stock concentration was determined using NanoDrop.

*In-vitro Droplets Assays via Confocal Microscopy*

Recombinant proteins were diluted to various stock concentrations using 1X storage buffer without PEG (Polyethylene Glycol 8000). For NaCl titration assays, the original protein stock (with 125 mM NaCl) was either diluted using 1X storage buffer with no NaCl or mixed with 1X storage buffer containing 2M NaCl to achieve final NaCl concentration ranging from 50 mM to 500 mM before imaging. For imaging,10 µL of fluorescent protein-fused condensate proteins were mixed with 10 µL of 20% (m/v) PEG solution in a PCR tube and allowed to incubate at room temperature for 30 min. Next, 10 μL of the protein mixture was added into a homemade flow chamber consisting of a glass slide (Corning, 2947-75X25) sandwiched by a coverslip (VWR, 48366-045) with a single layer of double-sided tape serving as a spacer. Images were acquired using a Zeiss LSM 980 Microscopy System, which includes an inverted Axio Observer microscope and a multiplex Airyscan module. Lasers at 405 nm, 488 nm and 567 nm were used for excitation, and appropriate PMT detectors were used for emission. As a negative control, pre-formed condensate solutions were mixed with an equal volume of 20% (w/v) 1,6-Hexanediol for 10 min at room temperature before imaging. For the droplet mixing assay, a 20 μM NPM1-Halo protein solution was incubated separately with equimolar amounts of each dye (TMR-Cl, Coumarin-Cl, or R110-Cl) for 10 min at room temperature. The different dye-labelled NPM1-Halo proteins were then incubated with an equal volume of 20% (w/v) PEG separately for another 30 min. Subsequently, these colored NPM1-Halo condensates were combined into a single solution and placed in the flow chamber for imaging. For RNA induced condensate formation, 20 μM mCherry-NPM1 was mixed with 50 ng/μL RNA-AZDye488 (without PEG) at room temperature for 30 min before imaging.

*Turbidity Assay*

To further characterize condensate formation beyond confocal microscopy, we performed turbidity measurement using mCherry-NPM1 protein. Briefly, condensate formation was assessed by mixing mCherry-NPM1 solutions with 1X storage buffer, followed by incubation with 20% (w/v) PEG for 30 min at room temperature. The resulting condensate solutions were transferred to an opaque 384-well-plate (Millipore Sigma, MZHVN0W50) in triplicate, and turbidity was measured at 600 nm using a microplate reader (Agilent, BioTek Synergy H1). The turbidity of mCherry-NPM1/PEG mixtures was analyzed across a range of mCherry-NPM1 concentrations, from 0.1 μM to 1 mM. To evaluate condensate growth over time, we prepared a mixture of 20 μM mCherry-NPM1 with 10% (w/v) PEG and monitored turbidity changes over a 30-minute time course.

*Condensate Imaging for Colocalization Assays*

To assess the interaction and exchange properties of protein condensates with macromolecules in the surrounding solution, colocalization assays were performed using both confocal microscopy and flow cytometry. Colocalization was evaluated in three categories: (a) antibodies, (b) lipids, and (c) cancer drugs. For antibody-based assays, a positive control was established using a 6xHisTag monoclonal antibody conjugated to Alexa Fluor 488 (Thermo Fisher, MAI-21315-A488).

A biotin monoclonal antibody labeled with Alexa Fluor 488 (Thermo Fisher, 53-9895-82) was used as a negative control. Recombinant mCherry-NPM1 protein, containing a 6xHisTag, was incubated with either anti-HisTag (1:800 dilution) or anti-biotin antibody (1:800 dilution) for 10 min at room temperature. The antibody-protein mixture was then incubated with PEG to facilitate condensate formation for 30 min, after which the samples were imaged. For lipid-based colocalization, 2 μM of Oregon Green 488 carboxylic acid (AAT Bioquest, 5-OG488) or Oregon Green phosphatidylethanolamine (Thermo Fisher, O12650) was incubated with 20 μM mCherry-NPM1 at room temperature for 10 min. The lipid-protein mixture was then combined with PEG for an additional 30-minute incubation to allow condensate formation. As an additional control, 20 μM mCherry-NPM1 was pre-incubated with 2 mg/mL neomycin trisulfate salt hydrate (Millipore Sigma, N1876) for 10 min, followed by incubation with 2 μM Oregon Green phosphatidylethanolamine for 10 min. PEG was then added for another 30-minute incubation prior to imaging. For cancer drug-based assays, 20 μM Coumarin-labeled NPM1-Halo protein was first incubated with PEG to form condensates. Following this, the samples were incubated with either 50 μM mitoxantrone (AmBeed, A207951) or 100 μM FLXT1 (MedChem Express, HY-119437) for 10 min at room temperature. The samples were subsequently pipetted for imaging.

*Condensate Analysis by Flow Cytometry*

To assess condensate formation and dynamics, flow cytometry was performed in parallel with confocal imaging using the same batch of protein samples, all analyzed on the same day as described above. Briefly, 35 μL of each 40 μM scaffolding protein (mCherry-NPM1, mCherry-DDX4, mCherry-HP1α, or EGFP-HMGB1) was mixed with 35 μL of either 1X storage buffer or 20% (w/v) PEG in a 96-well clear, untreated round-bottom plate (VWR, 82050-622). The mixtures were incubated at room temperature for 30 minutes, after which samples were analyzed on an Attune™ NxT Acoustic Focusing Cytometer. For the dissolution assay, pre-formed 20 μM condensates were treated with 10% (w/v) 1,6-hexanediol for an additional 10 minutes prior to flow cytometry analysis. Each condition was tested in triplicate, and the mean fluorescence intensity from over 10,000 events was used for quantification. To evaluate concentration-dependent condensate formation, varying concentrations of mCherry-NPM1 (2 -80 μM) were prepared by mixing protein stock solutions with either 1X storage buffer or 20% (w/v) PEG. Samples were incubated for 30 minutes at room temperature before flow cytometry analysis. For the salt titration experiments, mCherry-NPM1 was diluted in 1x storage buffer supplemented with either 0 or 2 M NaCl, yielding final NaCl concentrations ranging from 100 mM to 1 M and a constant final protein concentration of 20 µM. Equal volumes of 20% (w/v) PEG were added, and condensates were allowed to form for 30 minutes before analysis. For colocalization assays, antibodies, lipids, and small-molecule drugs were added to mCherry-NPM1 condensates at the same concentrations used in confocal microscopy experiments. These samples were similarly incubated and analyzed by flow cytometry.

To investigate condensate dynamics, a time-resolved assay was performed using NPM1-Halo. Unlike the high-throughput format, these experiments were carried out in individual tubes. Specifically, 10 µM NPM1-Halo-TAMRA was incubated with 10% (w/v) PEG for 1 to 12 hours at room temperature to allow condensate formation and aging. A baseline measurement was collected at - 5 minutes, and soluble NPM1-Halo-Coumarin (without PEG) was added to the pre-aged NPM1-Halo-TAMRA condensates, followed by immediate vortexing for 2 seconds. The samples were then monitored by flow cytometry at defined time intervals to track incorporation of the coumarin-labeled protein over time.

*Imaging Flow Cytometry*

To directly visualize protein condensates and compare their morphological features across different concentrations, imaging flow cytometry (IFC) was performed using the Amnis ImageStreamX Mark II system (Luminex Corporation). mCherry-NPM1 protein samples were prepared at concentrations ranging from 1 µM to 40 µM and mixed with an equal volume of 20% (w/v) PEG in a 96-well plate. Following a 30-minute incubation at room temperature to allow condensate formation, samples were subjected to IFC analysis. The instrument was equipped with a 60X objective and a 561 nm excitation laser for mCherry detection. Brightfield and fluorescent images of individual condensates were acquired simultaneously. A minimum of 10,000 events were collected per condition, and IDEAS® software (version 6.2, Luminex Corporation) was used for image analysis, including quantitative extraction of morphological parameters (e.g., area, circularity) and fluorescence intensity. Representative images of single mCherry-NPM1 condensates were selected across the concentration range to illustrate size and intensity changes. Surface area measurements derived from brightfield images were plotted against fluorescence intensity from the mCherry channel to assess concentration-dependent condensate growth. The combination of high-content imaging and quantitative flow analysis confirmed that condensates remain intact under flow and are amenable to both morphological and fluorescence-based characterization.

*Data Processing and Statistical Analyses*

All statistical analyses were performed using GraphPad Prism 9.5 (GraphPad Software). Bar graphs represent the mean ± standard deviation (SD) unless otherwise noted.

For conventional flow cytometry, over 10,000 events were collected per sample using the Attune™ NxT Acoustic Focusing Cytometer in volumetric mode (40 μL total acquisition volume at 200 μL/min). No gating was applied; all events were included in the analysis. The X-mean fluorescence intensity was used for quantification and plotting. Forward and side scatter (FSC/SSC) were recorded to monitor event quality, but not used for gating.

For imaging flow cytometry (IFC), >10,000 individual condensates were analyzed using the Amnis ImageStreamX Mark II system with IDEAS® software. Representative condensate images and quantitative data—such as surface area and mCherry fluorescence intensity—were extracted and displayed using scatter plots and violin plots. Condensates were identified using automated segmentation masks without manual gating or filtering.

For condensate dynamic exchange assays, quadrant analysis was used to monitor the incorporation of incoming blue-labeled proteins into pre-formed red-labeled condensates. Events were plotted based on red (TAMRA) and blue (coumarin) fluorescence, and the percentage of double-positive condensates was used to quantify molecular exchange over time. Flow cytometry settings were optimized to detect biomolecular condensates and are as follows: Acquisition volume: 40 μL (200 μL/min); Stopping option: 10,000 events; Laser voltages: FSC = 100, SSC = 240, BL1 = 220, YL1 = 230, YL2 = 230, all others = 200; Threshold: SSC = 500 ($0.5 \times 10^3$).

*Coarse-Grained (CG) Simulation Details*

We employed coarse-grained (CG) molecular dynamics simulations to investigate the phase behavior and intermolecular interactions of full-length NPM1 proteins. Each protein was modeled using a $C_\alpha$-level representation, wherein every amino acid residue is mapped to a single bead centered on its $\alpha$-carbon ($C_\alpha$) position. The interactions between residues were described by the HPS-Urry model,[4] which has been shown to accurately reproduce liquid-liquid phase separation (LLPS) in silico.[5, 6] To account for the structural organization of NPM1, we applied rigid-body motion to its folded domains—namely, the N-terminal oligomerization domain (OD; residues 14–118) and the C-terminal DNA-binding domain (CTD; residues 243–294), based on high-confidence structure predictions (pLDDT > 90). The remaining segments, including the N-terminal residues (1–13) and the central intrinsically disordered region (IDR; residues 119–242), were treated as fully flexible.

Simulations were conducted in a rectangular slab geometry of dimensions $175 \times 175 \times 1225$ Å$^3$ containing 100 NPM1 chains, with periodic boundary conditions applied in all three directions. We performed a $5\ \mu s$ long Langevin dynamics simulations at a fixed temperature of 300 K, with the friction coefficient $\gamma = m_{AA}/\tau$. Here $m_{AA}$ is the mass of each amino acid bead and $\tau$ is the damping factor set to 1000 ps. The equations of motion were integrated using a velocity-Verlet algorithm with a time step of 10 fs. All simulations were performed using the HOOMD-blue molecular dynamics engine (version 4.7) with additional features provided by the azplugins package.[7] We conducted the CG simulation at 100 mM salt concentration.

For analysis, we excluded the initial $1\ \mu s$ of the trajectory to allow for equilibration. Density profiles and intermolecular contact maps were computed from the remaining $4\ \mu s$ of production data. Error bars were calculated using block averaging by dividing the production trajectory into 4 equal blocks and computing the standard error of the mean.

*Theoretical Details of First-Passage Calculations*

Let us define $F_{dil}(t)$ as the first-passage probability density function of entering a blue protein into the dense phase of the red condensate for the first time at time $t$ if initially at $t = 0$, the system started in the dilute phase. Similarly, one can define the first-passage probability density function for the interphase as $F_{int}(t)$. Then the temporal evolution of these probability functions is governed by a set of backward master equations[8]:

$$\frac{F_{dil}(t)}{dt} = k_{on}F_{int}(t) - k_{on}F_{dil}(t) \quad \text{(S1)}$$

$$\frac{F_{int}(t)}{dt} = k_{bounce}F_{dil}(t) + k_{entry}F_{den}(t) - (k_{bounce} + k_{entry})F_{int}(t) \quad \text{(S2)}$$

In this equation, $F_{den}(t)$ is the probability density of a blue protein to be found in the dense phase immediately after leaving the interphase, and we can assume that $F_{den}(t) = \delta(t)$. This means that if the system is in this state at $t = 0$, the process is immediately accomplished.

Applying Laplace transformations $\tilde{F}(s) \equiv \int_0^\infty e^{-st} F(t)dt$, we obtain

$$(s + k_{on})\widetilde{F_{dil}}(s) = k_{on}\widetilde{F_{int}}(s) \quad \text{(S3)}$$

$$(s + k_{bounce} + k_{entry})\widetilde{F_{int}}(s) = k_{bounce}\widetilde{F_{dil}}(s) + k_{entry} \tag{S4}$$

Solving Eqs. (S3 – S4) simultaneously, we obtain

$$\widetilde{F_{dil}}(s) = \frac{k_{on}k_{entry}}{s\big(s + k_{on} + k_{bounce} + k_{entry}\big) + k_{on}k_{entry}} \tag{S5}$$

After performing inverse Laplace transformation of the expression (S5), we obtain the first-passage distribution function of exchange times from dilute to dense phase and it is given by

$$F(t) \equiv F_{dil}(t)$$
$$= \frac{e^{-\frac{t}{2}\left(k_{on}+k_{bounce}+k_{entry}+\sqrt{\left(k_{on}+k_{bounce}+k_{entry}\right)^2-4k_{on}k_{entry}}\right)}\left(e^{t\sqrt{\left(k_{on}+k_{bounce}+k_{entry}\right)^2-4k_{on}k_{entry}}} - 1\right)k_{on}k_{entry}}{\sqrt{(k_{on} + k_{bounce} + k_{entry})^2 - 4k_{on}k_{entry}}} \tag{S6}$$

Then we can compute the cumulative distribution function (CDF) from this first-passage time distribution:

$$CDF(t) = \int_0^t F(\tau)d\tau \tag{S7}$$

Finally, the mean first-passage times (MFPT) of protein exchange from dilute to dense phase can be obtained as

$$T_{exchange} = \int_0^\infty tF_{dil}(t)dt = -\frac{\partial\widetilde{F_{dil}}(s)}{\partial s}|_{s=0} = \frac{k_{on} + k_{bounce} + k_{entry}}{k_{on}k_{entry}} \tag{S8}$$

**Table S1. Corresponding protein sequences**

**1) pET-28a-HaloTag-NPM1**

>Amino acid sequence

MHHHHHHGIEENLYFQSGSGMAEIGTGFPFDPHYVEVLGERMHYVDVGPRDGTPVLFLHGNP
TSSYVWRNIIPHVAPTHRCIAPDLIGMGKSDKPDLGYFFDDHVRFMDAFIEALGLEEVVLVIHDW
GSALGFHWAKRNPERVKGIAFMEFIRPIPTWDEWPEFARETFQAFRTTDVGRKLIIDQNVFIEGT
LPMGVVRPLTEVEMDHYREPFLNPVDREPLWRFPNELPIAGEPANIVALVEEYMDWLHQSPVP
KLLFWGTPGVLIPPAEAARLAKSLPNCKAVDIGPGLNLLQEDNPDLIGSEIARWLSTLEISGGSG
MYTDMEDSMDMDMSPLRPQNYLFGCELKADKDYHFKVDNDENEHQLSLRTVSLGAGAKDEL
HIVEAEAMNYEGSPIKVTLATLKMSVQPTVSLGGFEITPPVVLRLKCGSGPVHISGQHLVAVEED
AESEDEDEEDVKLLGMSGKRSAPGGGNKVPQKKVKLDEDDEDDDEDDEDDEDDDDDDFDEE
ETEEKVPVKKSVRDTPAKNAQKSNQNGKDLKPSTPRSKGQESFKKQEKTPKTPKGPSSVEDIK
AKMQASIEKGGSLPKVEAKFINYVKNCFRMTDQEAIQDLWQWRKSLV

>DNA sequence

ATGCACCATCACCATCACCATGGGATCGAGGAAAACCTGTACTTCCAATCCGGTTCTGGAAT
GGCCGAAATTGGCACCGGCTTTCCGTTTGATCCGCATTATGTTGAAGTCCTGGGCGAACGC
ATGCACTATGTGGATGTGGGTCCGCGTGATGGTACCCCGGTCCTGTTTCTGCATGGCAACC
CGACGAGCAGCTATGTTTGGCGCAATATTATCCCGCATGTTGCACCGACCCACCGTTGCATT
GCCCCGGATCTGATCGGCATGGGCAAAAGCGATAAACCGGATCTGGGCTATTTCTTTGATG
ATCACGTGCGCTTTATGGATGCGTTTATTGAAGCCCTGGGCCTGGAAGAAGTGGTTCTGGT
TATCCATGATTGGGGCAGCGCACTGGGTTTTCACTGGGCCAAACGCAACCCGGAACGTGTT
AAAGGCATTGCGTTTATGGAATTTATTCGCCCGATCCCGACCTGGGATGAATGGCCGGAATT
TGCCCGTGAAACGTTTCAGGCGTTTCGCACCACGGATGTGGGCCGTAAACTGATCATCGAT
CAGAACGTTTTCATCGAGGGTACGCTGCCGATGGGCGTCGTGCGTCCGCTGACGGAAGTT
GAAATGGATCATTATCGTGAACCGTTTCTGAATCCGGTCGATCGCGAACCGCTGTGGCGTTT
TCCGAACGAACTGCCGATTGCGGGCGAACCGGCCAATATCGTCGCGCTGGTTGAAGAATAT
ATGGATTGGCTGCACCAGAGCCCGGTCCCGAAACTGCTGTTTTGGGGTACCCCGGGCGTG
CTGATTCCGCCGGCCGAAGCGGCCCGCCTGGCGAAAAGCCTGCCGAATTGTAAAGCCGTG
GATATCGGCCCGGGCCTGAACCTGCTGCAGGAAGATAATCCGGATCTGATTGGCAGCGAAA
TCGCGCGTTGGCTGAGCACGCTGGAAATCAGCGGCGGTTCTGGAATGTACACGGATATGG
AAGACTCGATGGATATGGACATGAGTCCTCTTAGGCCTCAGAACTACCTTTTCGGCTGTGAA
CTAAAGGCTGACAAAGACTATCACTTTAAAGTGGATAATGATGAAAATGAGCACCAGTTGTCA
TTAAGAACGGTCAGTTTAGGAGCAGGGGCAAAAGATGAGTTACACATCGTAGAGGCAGAAG
CAATGAACTATGAAGGCAGTCCAATTAAAGTAACACTGGCAACTTTGAAAATGTCTGTACAAC
CAACAGTTTCCCTAGGGGGCTTTGAAATTACACCACCTGTGGTCTTACGGTTGAAGTGTGG
TTCAGGGCCTGTGCACATTAGTGGACAGCATCTAGTAGCTGTAGAGGAAGATGCAGAGTCT
GAAGATGAAGATGAGGAGGACGTAAAACTCTTAGGCATGTCTGGAAAGCGATCTGCTCCTG
GAGGTGGTAACAAGGTTCCACAGAAAAAAGTAAAACTTGATGAAGATGATGAGGACGATGAT
GAGGACGATGAGGATGATGAGGATGATGATGATGATGATTTTGATGAAGAGGAAACTGAAGA
AAAGGTCCCAGTGAAGAAATCTGTACGAGATACCCCAGCCAAAAATGCACAAAAATCAAACC
AAAATGGAAAAGACTTAAAACCATCAACACCGAGATCAAAGGGTCAAGAGTCCTTCAAAAAA
CAGGAAAAGACTCCTAAAACACCAAAAGGACCTAGTTCTGTAGAAGACATTAAGGCAAAAAT
GCAAGCAAGTATAGAAAAAGGCGGTTCTCTTCCCAAAGTGGAAGCCAAGTTCATTAATTATG

TGAAGAATTGTTTCCGGATGACTGACCAGGAGGCTATTCAAGATCTCTGGCAGTGGAGGAA
ATCTCTTGTTTAA
